# WASP integrates substrate topology and cell polarity to guide neutrophil migration

**DOI:** 10.1101/2021.05.12.443722

**Authors:** Rachel M. Brunetti, Gabriele Kockelkoren, Preethi Raghavan, George R. R. Bell, Derek Britain, Natasha Puri, Sean R. Collins, Manuel D. Leonetti, Dimitrios Stamou, Orion D. Weiner

## Abstract

To control their shape and movement, cells leverage nucleation promoting factors (NPFs) to regulate when and where they polymerize actin. Here we investigate the role of the immune-specific NPF WASP during neutrophil migration. Endogenously-tagged WASP localizes to substrate-induced plasma membrane deformations. Super-resolution imaging of live cells reveals that WASP preferentially enriches to the necks of these substrate-induced membrane invaginations, a distribution that could support substrate pinching. Unlike other curvature-sensitive proteins, WASP only enriches to membrane deformations at the cell front, where it controls Arp2/3 complex recruitment and actin polymerization. Despite relatively normal migration on flat substrates, WASP depletion causes defects in topology sensing and directed migration on textured substrates. WASP therefore both responds to and reinforces cell polarity during migration. Surprisingly, front-biased WASP puncta continue to form in the absence of Cdc42. We propose that WASP integrates substrate topology with cell polarity for 3D guidance by selectively polymerizing actin around substrate-induced membrane deformations at the leading edge. A misregulation of WASP-mediated contact guidance could provide insight into the immune disorder Wiskott-Aldrich syndrome.

## Introduction

Motile cells must coordinate many types of actin networks to achieve directed migration (Blanchoin et al., 2014). Nucleation promoting factors (NPFs) control when and where branched actin is assembled, with different NPFs giving rise to different types of actin networks (Pollitt and Insall, 2009; Rottner et al., 2017). For example, WAVE forms broad, propagating waves at the leading edge that pattern the sheet-like actin networks that generate lamellipodia (Weiner et al., 2007; Veltman et al., 2012; Leithner et al., 2016). In contrast, N-WASP has a punctate distribution and forms finger-like actin networks that comprise podosomes (Isaac et al., 2010; Nusblat et al., 2011; Mizutani et al., 2002), invadopodia (Yamaguchi et al., 2005; Yu et al., 2012; Yu and Machesky, 2012), and sites of endocytosis (Kessels and Qualmann, 2002; Merrifield et al., 2004; Benesch et al., 2005). While the function of WAVE and N-WASP is well understood, other NPFs remain poorly characterized. This includes WASP, an N-WASP homologue specific to the hematopoietic lineage. WASP was initially identified through its role in Wiskott-Aldrich syndrome (Derry et al., 1994), an immune disorder characterized by abnormal platelets, eczema, and recurrent infection (Wiskott, 1937; Aldrich et al., 1954). However, despite the homing defects of immune cells in WASP-deficient patients and animal models (Ochs et al., 1980; Snapper et al., 2005; De Noronha et al., 2005; Westerberg et al., 2005; Jones et al., 2013), how WASP contributes to actin organization and cell migration remains poorly understood.

The function of WASP in vertebrates has been inferred from the function of its homologue N-WASP. However, important functional divergences have been reported between these proteins (Isaac et al., 2010; Nusblat et al., 2011). In particular, N-WASP cannot compensate for the loss of WASP in T cell chemotaxis (Jain and Thanabalu, 2015). The basis of WASP’s role in immune cell guidance remains unknown. Do functions ascribed to N-WASP such as endocytosis and podosome/invadopodia formation also underlie WASP-deficient immune cell migration defects? Or does WASP participate in additional aspects of physiology during immune cell migration? The functional roles of actin nucleators and NPFs have successfully been elucidated through analysis of spatiotemporal dynamics and loss-of-function phenotypes (Bear et al., 1998; Rogers et al., 2003; Kunda et al., 2003; Weiner et al., 2007; Miki et al., 1998; Nakagawa et al., 2001; Taunton et al., 2000; Benesch et al., 2002; Duleh and Welch, 2010; Zuchero et al., 2009; Sagot et al., 2002; Yang et al., 2007; Manor et al., 2015). Here we use this approach to probe WASP’s contribution to neutrophil migration.

By fluorescently labeling endogenous WASP, we find its primary localization to be focal puncta on the ventral surface of cells. These structures are not sites of clathrin-mediated endocytosis but are instead triggered by interaction with the substrate. Specifically, sites where the substrate induces inward plasma membrane curvature lead to local WASP enrichment. Super-resolution imaging reveals that WASP concentrates at the necks of substrate-induced invaginations, a distribution that supports engagement with substrate features. Typically, curvature-responsive proteins show the same degree of curvature sensitivity in different parts of the same cell (Zhao et al., 2017; Lou et al., 2019). In contrast, WASP integrates cell polarity with membrane curvature sensing and enriches at sites of inward curvature only in the front half of the cell. This spatial restriction could enable the cell to disengage with the substrate at the trailing edge while prioritizing engagement in the direction of advance. Through analyzing knockouts, we find that WASP is essential for Arp2/3 complex recruitment and actin polymerization at sites of membrane deformation at the cell front, linking the substrate’s topology with the migration machinery. Consistent with this idea, WASP-null cells exhibit migration defects on patterned substrates, while their motility on flat substrates is relatively unaffected. Our data establishes a role for WASP in connecting substrate topology and cell polarity with migration in neutrophils.

## Results

### WASP enriches to puncta that associate with the substrate

Because of its key role in the immunological disorder Wiskott-Aldrich Syndrome, WASP has traditionally been studied through its loss-of-function phenotypes in immune cells from patients and animal models of the disease (Ochs et al., 1980; Snapper et al., 2005; De Noronha et al., 2005; Westerberg et al., 2005; Jones et al., 2013). Less attention has been paid to its localization and dynamics in cells. Therefore, we turned to the easily manipulated human neutrophil-like cell line HL-60 to better understand the spatiotemporal dynamics of WASP and how these attributes might inform its function. In a previous report, exogenously expressed WASP was shown to enrich to the tip of the leading edge and to membrane-associated puncta in HL-60s (Fritz-Laylin et al., 2017). However, protein overexpression can lead to ectopic localization (Doyon et al., 2011). To better profile WASP’s functional properties, we generated a fluorescently tagged protein at the endogenous locus in HL-60 cells using CRISPR-Cas9 and homology-directed repair (Fig. S1A-C).

WASP knock-in HL-60 cells were confined in 2D to flatten the ventral surface of the cell against the coverslip and facilitate imaging by TIRF microscopy. Endogenous WASP localized to membrane-associated puncta and the periphery of the leading edge, a distribution which was broadly consistent with exogenous WASP (Fritz-Laylin et al., 2017). However, enrichment of WASP to puncta in the front half of the cell, separate from the cell periphery, was more strongly pronounced than previously appreciated and proved to be the dominant pattern of WASP localization (Fig. 1A and 1C; Video 1). Additionally, WASP puncta were remarkably stationary, often persisting at the same position relative to the coverslip on the minute time scale despite bulk cell displacement (Fig. 1B). Because N-WASP has been implicated in clathrin-mediated endocytosis (CME) (Kessels and Qualmann, 2002; Merrifield et al., 2004; Benesch et al., 2005), we investigated whether WASP puncta represent sites of CME. Endogenous WASP and endogenous clathrin light chain A failed to co-localize at the cell front (Fig. 1D-E;Fig. S1D-E; Video 2). In addition, both clathrin-dependent and independent endocytic pathways have been shown to occur at the rear of polarized HL-60 cells (Davis et al., 1982; Subramanian et al., 2018). Therefore, front-biased WASP puncta (Fig. 1E) do not appear to mark sites of endocytosis.

**Fig. 1.**
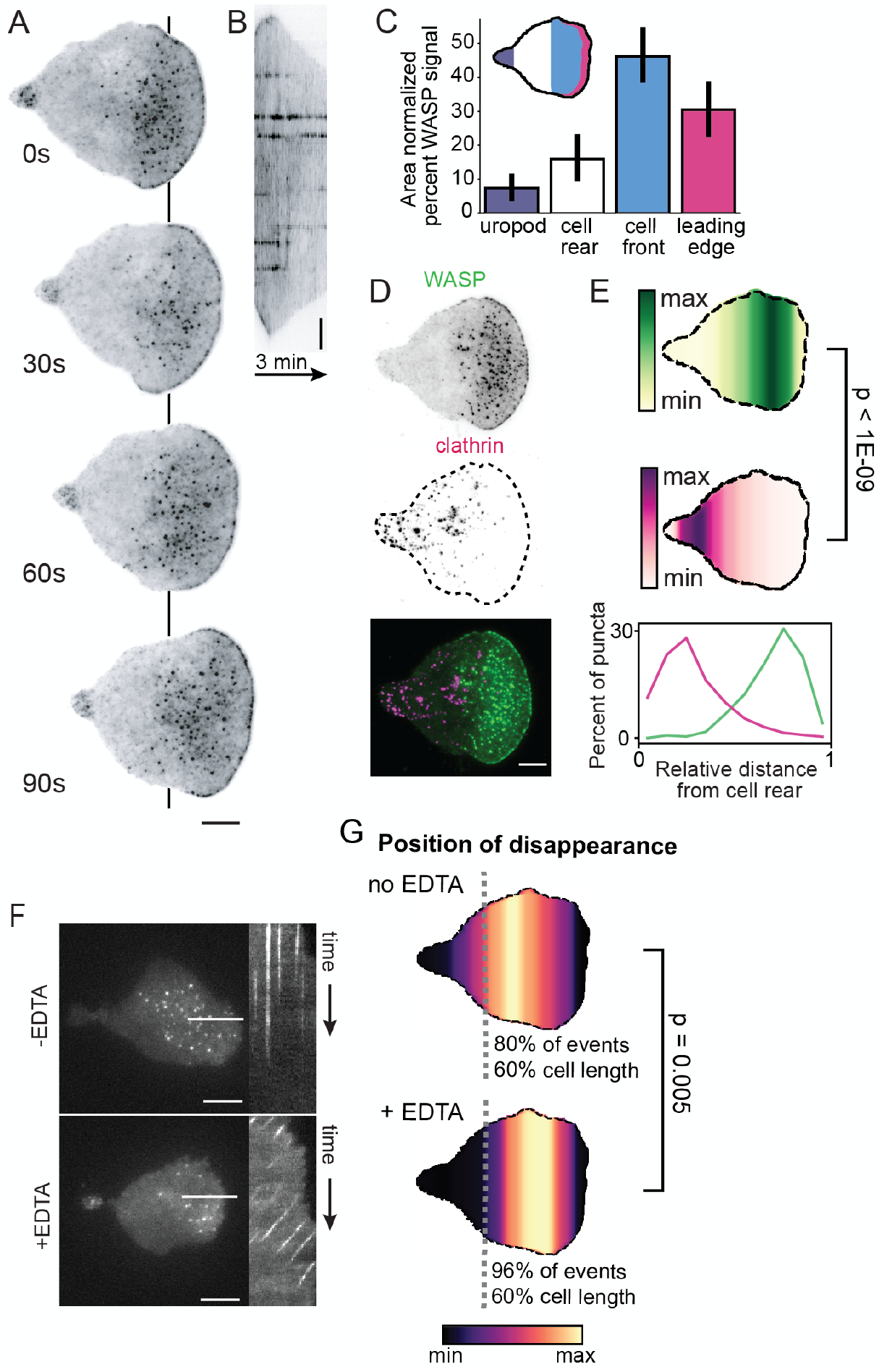
Endogenous WASP forms puncta stabilized by cell-substrate interactions. (A) TIRF microscopy of endogenous WASP reveals localization to the tip of the lamellipod and to ventral puncta throughout the front half of the cell and to a lesser extent the uropod. Color map is inverted. Each frame is 30s. See also Video 1. (B) Kymograph along the line in (A) highlights the stationary nature of WASP puncta relative to the substrate. Scale bar is 2 μm. (C) WASP shows primary localization to punctate structures biased toward the cell front. n = 22 cells collected across three experiments. Leading edge: 30.5 ± 4.4%, cell front: 46.1 ± 4.4%, cell rear: 16.0 ± 3.6%, uropod: 7.5 ± 2.2% of total WASP signal. Values are area normalized. (D) Endogenous WASP and endogenous clathrin light chain A fail to co-localize. Color map is inverted for single channel images. See also Video 2. (E) Spatial distribution of WASP (green) and clathrin (magenta) puncta and their corresponding line scans confirm consistent spatial separation of these proteins in co-expressing cells. n = 11 cells collected across three experiments with more than 1000 puncta measured for each marker. Mean relative position of WASP is 0.71 ± 0.01 and mean relative position of clathrin is 0.29 ± 0.02 on the single cell level. p= 8.82E-10 by a paired two-tailed t-test on the mean puncta position of each marker in each cell. (F) Treatment of cells with EDTA, a calcium and magnesium chelator that blocks integrin-based adhesion, causes normally stationary WASP puncta to experience retrograde flow relative to the substrate. Left depicts a single frame of endogenous WASP with a line used to construct the kymograph to the right. See also Video 3. (G) Spatial distribution of the position of disappearance of WASP puncta reveals earlier extinction in the presence of EDTA. n = 12 cells per condition collected across three experiments with approximately 300 puncta measured for each condition. Mean relative position of puncta disappearance is 0.55 ± 0.03 in the absence of EDTA and 0.66 ± 0.02 in the presence of EDTA on the single cell level. p = 0.005 by an unpaired two-tailed t-test on the mean position of puncta disappearance for each cell. For relative positions, the cell rear is defined as 0 and the cell front is defined as 1. Scale bars are 5 μm unless otherwise specified.

Other documented roles for WASP/N-WASP include the formation of invasive structures (Yu et al., 2012; Yu and Machesky, 2012; Isaac et al., 2010; Nusblat et al., 2011; Mizutani et al., 2002; Yamaguchi et al., 2005) and facilitation of cell adhesion (Misra et al., 2007; Zhang et al., 2006) and spreading (Misra et al., 2007; Liu et al., 2013). Neutrophils have a limited capacity for generating invadopodia or podosomes, failing to invade soft substrates like matrigel or to form their characteristic actin-vinculin rosettes (Cougoule et al., 2012). We therefore investigated whether WASP puncta are linked to cell adhesion. While neutrophils do not generate the long-lived focal adhesions found in many adherent cell lines (Yuruker and Niggli, 1992; Lämmermann and Sixt, 2009), we hypothesized that the fixed location of WASP puncta relate to adhesion with the underlying substrate. To test this, we confined cells between two glass coverslips and treated them with EDTA, a calcium and magnesium chelator that blocks integrin-ligand binding and therefore prevents adhesion (Zhang and Chen, 2012). Under inelastic confinement, neutrophils can migrate in an adhesion-independent manner (Malawista et al., 2000), using friction to chimney between the glass surfaces like a climber squeezing between two rock faces. In the absence of integrin-based adhesion, we found that WASP puncta continued to form on the cell surface but were no longer stationary relative to the substrate and instead began undergoing retrograde flow (Fig. 1F; Video 3). Additionally, WASP puncta are less stable for cells lacking integrin adhesions, extinguishing significantly closer to their point of nucleation at the cell front (Fig. 1G). The induced motility of WASP puncta when cell-substrate adhesions are removed suggests that stationary WASP puncta normally depend on engagement with the substrate. This finding is supported by previous observations that WASP-null neutrophils have an impaired ability to adhere to and cross cell monolayers under shear flow (Zhang et al., 2006) while neutrophils expressing only constitutively active WASP demonstrate increased adhesion and migration in flow chambers (Keszei et al., 2018).

Based on these data, we hypothesized that WASP puncta couple the lamellipodial actin network to the cell substrate to aid in motility. We next asked which substrate features are involved in WASP recruitment.

### WASP is curvature-sensitive and sorts to sites of saddle curvature

Positive (inward) membrane curvature can organize N-WASP both in reconstituted systems (Takano et al., 2008) and in live cells plated on nanopatterns (Lou et al., 2019). We sought to test whether inward curvature similarly templates WASP organization. However, neutrophils only loosely adhere to their substrate (Lämmermann and Sixt, 2009), limiting; the ability of nanopatterns to precisely deform the cell membrane. We therefore forced cells into closer apposition to the substrate by overlaying them with agarose and incubating samples until cells were flattened (Fig. 2A). Because WASP/N-WASP natively associates with spherical membrane invaginations like endocytic pits, we used polystyrene beads as our membrane-deforming agents. The use of spherical beads enabled us to explore 2D curvatures that are inaccessible with most fabricated nanopatterns. Specifically, beads allowed us to probe (1) two dimensions of positive (inward) isotropic curvature at the body of the bead-induced invagination and (2) saddle curvature at the invagination neck. Because curvature sensitive proteins can discriminate between 1D and 2D curvatures (Iversen et al., 2015; Larsen et al., 2020), it is important to have tools that can probe 2D curvature in cells. Additionally, this approach has the benefit of being highly tunable through the availability of beads of different diameters (and therefore radii of curvature). Finally, use of commercially available, inert beads removed the cost and throughput limitations of nanofabrication techniques like electron beam lithography (Li et al., 2019).

**Fig. 2.**
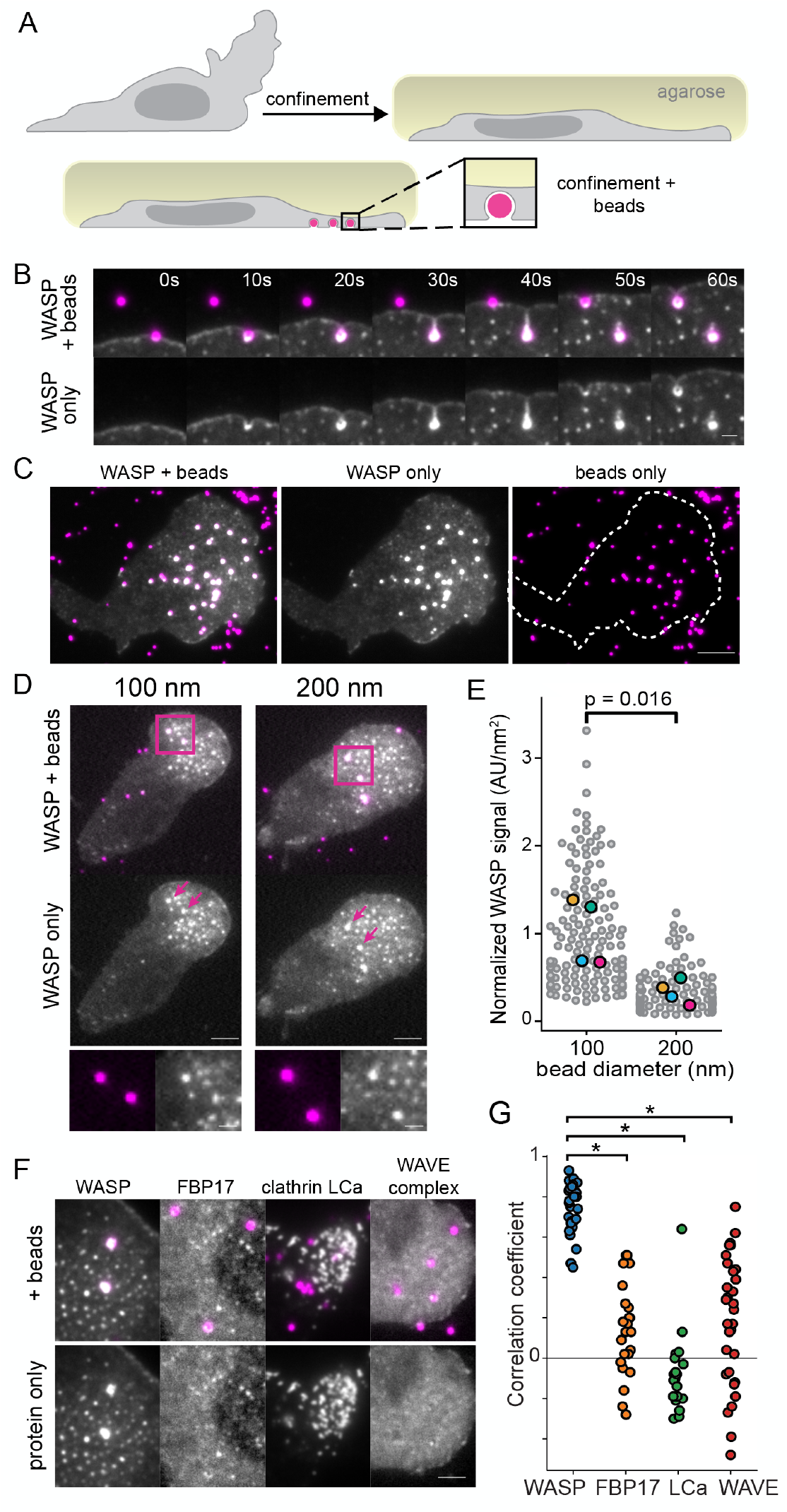
WASP is recruited to membrane invaginations in a curvature-dependent manner. (A) Experimental strategy of confining cells onto beads to induce sites of substrate-controlled plasma membrane curvature. (B) Time lapse imaging reveals rapid recruitment of endogenous WASP to 500 nm bead-induced invaginations. Scale bar is 1 μm. (C) WASP consistently localizes to 200 nm bead-induced invaginations across the cell front. Scale bar is 5 μm. See also Video 4. (D) WASP knock-in cells were confined onto beads of varying diameter to assess the curvature sensitivity of WASP recruitment. Images are scaled to the same intensity for WASP Below is a zoomed inset of the boxed regions. Scale bars are 5 μm and 1 μm for the inset. (E) Quantification of WASP signal at 100 and 200 nm beads reveals significantly more WASP at smaller invaginations (sites of higher curvature) when normalized to bead surface area. Mean WASP signal per unit area is 1.01 ±0.19 AU/nm^2^ for 100 nm beads and 0.34 ± 0.07 AU/nm^2^ for 200 nm beads on the replicate level. Large markers denote the mean of each replicate, which were compared using an unpaired two-tailed t-test: p = 0.016. Measurements are from 142 100 nm beads and 120 200 nm beads collected across four experiments. (F) WASP enrichment to 500 nm beads is much stronger than that of other known curvature-sensitive proteins: FBP17, clathrin light chain A (LCa), and WAVE complex. Scale bar is 2 μm. (G) Pearson correlation coefficients for 1.5X1.5 μm ROIs of a bead and the protein of interest. WASP has a significantly higher correlation with bead signal (r = 0.75 ± 0.02) compared to FBP17 (0.14 ± 0.05), clathrin LCa (−0.09 ± 0.04), and WAVE complex (0.19 ± 0.06). The correlation of each protein with beads was compared to the correlation of WASP with beads using an unpaired two-tailed t-test. Asterisks mark significance. p_WASP/FBP17_ = 6.56E-18, p_WASP/clathrin_ = 4.60E-24, and p_WASP/WAVE_ = 5.58E-13. n_WASP_ =31, n_FBP17_ = 24, n_clathrin_ =21, and n_WAVE_ = 31 beads each from one experiment.

At HL-60s migrated over beads, WASP was recruited within seconds to all beads and persisted while they were under the lamellipod (Fig. 2B-C; Video 4). To determine whether this enrichment is curvature sensitive, we campared WASP recruitment in cells migrating over 100 and 200 nm beads (Fig. 2D). *in vitro*, enrichment of curvature-sensitive proteins to different sized liposomes is normalized per unit area (Bhatia et al., 2009). Applying this normalization scheme to our data, we found a three-fold enrichment of WASP to the smaller bead size (higher radius of curvature) (Fig. 2E). This finding is consistent with measurements for curvature-sensitive proteins, including BAR domain-containing proteins (Bhatia et al., 2009) and components of CME (Zhao et al., 2017), suggesting that WASP recruitment is sensitive to curvature. To probe whether membrane deformation could similarly be an organizing cue for physiological substrates, we plated cells on coverslips coated with a thin layer of fluorescent collagen fibers (Elkhatib et al., 2017). Even without agarose confinement, cells were deformed by collagen fibers, and endogenous WASP was recruited to f ell-fiber interfaces (Fig. S2).

WASP was strikingly consistent in its recruitment across multiple beads under the same lamellipod. In adherent cells plated on nanopatterns, curvature-sensitive proteins are normally heterogeneous in their degree of recruitment to adjacent nanobars (Lou et al., 2019; Zhao et al., 2017). This led us to compare the strength of WASP recruitment with that of other known curvature sensors through measuring enrichment to sites of bead-induced membrane deformation (Fig. 2F-G). First, we investigated the WASP/N-WASP-interacting, BAR domain-containing protein FBP17. FBP17 activates N-WASP in *vitro* (Takano et al., 2008) and has been posited as an organizer of N-WASP-dependent actin nucleation in endocytosis (Tsujita et al., 2006) and at the leading edge of polarized cells (Tsujita et al., 2015). In COS-1 cells FBP17 also exists as membrane-associated puncta (Tsujita et al., 2015) that are reminiscent of our WASP puncta. Despite co-localization of endogenous FBP17 and WASP at puncta biased toward the cell front (Fig. S3A-B), FBP17 was neither required for WASP puncta formation nor WASP recruitment to bead-induced membrane deformations since these behaviors persisted in FBP17 knockout cells (Fig. S3F-G). Furthermore, the degree of FBP17 enrichment to beads was significantly poorer than that of WASP (Fig. 2G). Similarly, endogenous clathrin light chain A (LCa), which enriches to bead-induced membrane invaginations in MDA-MB-231 cells (Elkhatib et al., 2017), exhibited less robust recruitment to beads than WASP in neutrophils (Fig. 2G). Finally, since WAVE complex exhibits a preference for saddle curvatures (Pipathsouk et al., 2019), we measured its enrichment to bead-induced invaginations. While WAVE complex transiently enriched to some beads, it failed to persistently enrich (Fig. 2G). WASP appears to be unusual among other curvature-sensitive proteins in its consistent marking of 500 nm-scale membrane invaginations in neutrophils.

While we found that curvature informs WASP recruitment, it remained unclear what type of membrane geometry was driving this process since bead-induced invaginations have both positive (inward) isotropic curvature (along the bead body) and saddle curvature (at the invagination neck). To elucidate WASP’s preferred curvature, we turned to live-cell 2D and 3D super-resolution stimulated emission depletion (STED) microscopy of the membrane at bead-induced invaginations. Using 3D STED, we achieved a lateral (x-y) resolution of 160 ± 6 nm and an axial (x-z) resolution of 160 ± 16 nm (Fig. S4A-E). We began with cells confined onto fluorescent 500 nm beads to clearly separate the isotropically positive invagination body from the saddle-shaped neck. We observed two different classes of membrane organization around 500 nm beads, each of which correlated to distinct patterns of WASP enrichment (Fig. 3A). In one set of cases, the membrane wrapped around a bead but failed to cinch around the bead bottom, forming an inverted “U” shape (∩). In this case, WASP lined the entirety of the bead-induced membrane invagination (Fig. 3A, top panel). In a second set of cases, the membrane formed a cinched neck under the bead, making an “omega” shape (Ω) (Fig. 3A, bottom panels). When this morphology was observed, WASP no longer surrounded the entire structure but was instead concentrated onto the sides of the invagination neck. At smaller, 200 nm bead-induced invaginations, we could not resolve any membrane necks, and WASP organization at these sites appeared to match that of “U” shaped invaginations (Fig. 3B). In contrast, others have reported saddle-enrichment of N-WASP (Kaplan et al., 2020) and the yeast orthologue Las17 (Mund et al., 2018) on this size scale using STORM to image CME in fixed cells. Finally, we compared WASP intensities at uniformly coated 200 and 500 nm bead-induced invaginations measured with 3D STED and again found an increased enrichment to beads with higher radii of curvature (Fig. S4F).

**Fig. 3.**
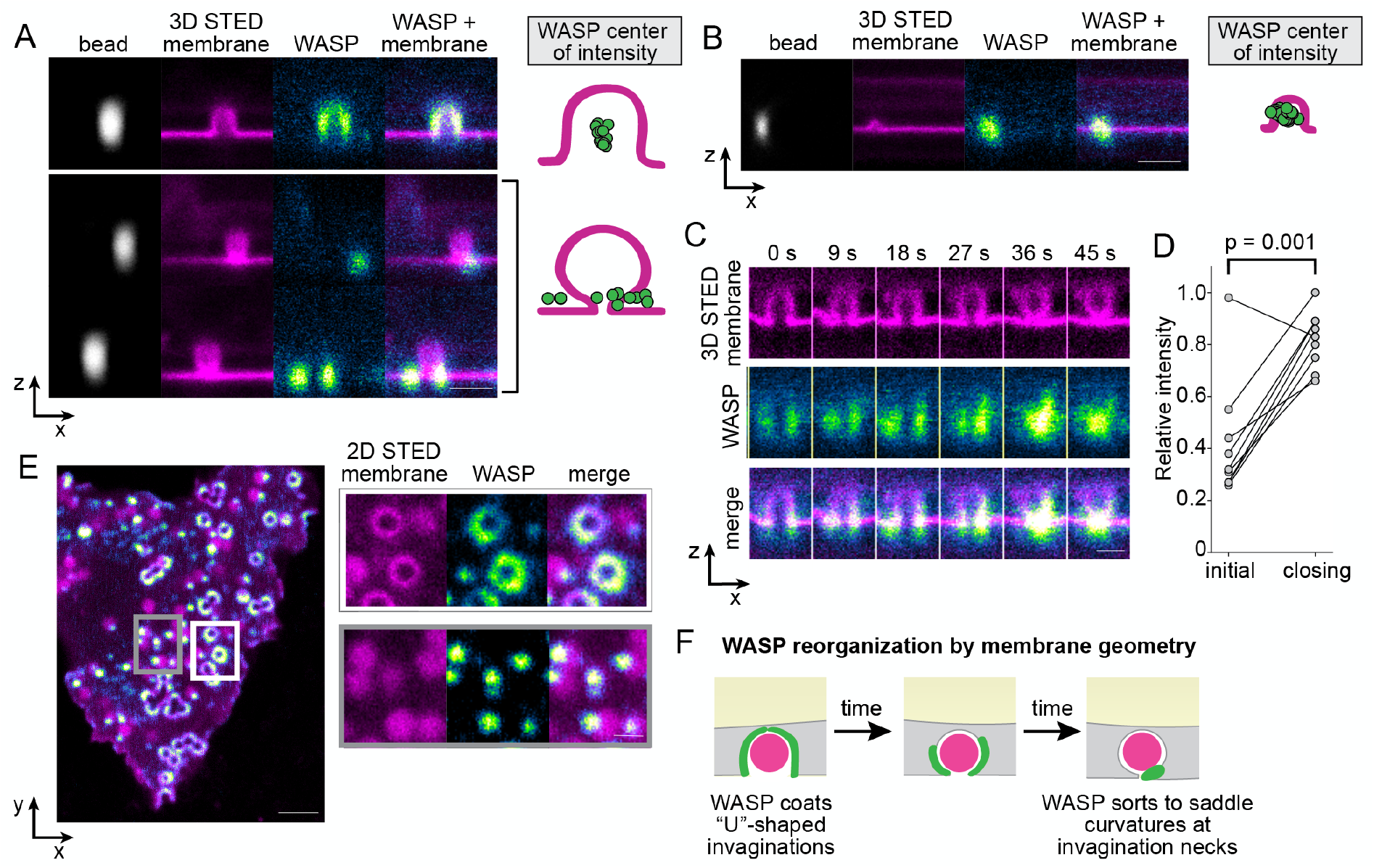
Super-resolution imaging reveals nanoscale reorganization of WASP by membrane geometry. (A) Live cell 3D STED microscopy of the membrane coupled with confocal imaging of WASP on 500 nm beads show that WASP enriches to the site of highest inward curvature, either across the whole bud (top) or at a neck when present (bottom). Right hand side depicts the center of intensity of WASP for these two cases, with n_500nm, no neck_ = 17 and n_500nm, neck_ = 11 invaginations collected across 8 and 4 cells, respectively, from the same experiment. Scale bar is 1 μm. (B) WASP enriches to the entire bead at smaller, 200 nm invaginations, as reflected in the center of intensity of WASP to the right. n_200nm_ = 29 invaginations collected across 9 cells from one experiment. Scale bar is 1 μm. (C) Time lapse 3D STED imaging of the membrane reveals evolution from an open to closed-neck structure at sites of bead-induced invagination. As this occurs, WASP redistributes and moves from the bead surface to the invagination neck. Scale bar is 500 nm. (D) As the membrane closes down around a bead, total WASP levels significantly increase. Mean normalized WASP intensity is 0.42 ± 0.08 AU at the first frame of the movie compared to 0.82 ± 0.04 AU at the time of neck closing on the single invagination level. A paired two-tailed t-test was used to compare the normalized intensity of WASP at these time points from nine invaginations collected across three experiments: p = 0.001. (E) 2D STED imaging of the cell membrane reveals both open and closed invaginations. At open invaginations WASP coats the base (white box). As invaginations close down, WASP is reorganized into a focal accumulation at the neck (gray box). Scale bars are 2 μm and 500 nm in the inset. See Video 5 for the dynamics of this process. (F) Model for WASP evolution as the membrane of a confined cell constricts around a bead.

We next investigated the temporal relation between the “U” and “omega” invagination states, and, using time-lapse 3D STED, were able to record “U”-shaped membranes closing around beads into “omega” shapes (Fig. 3C; Video 5). We found that not only does WASP reorganize during neck closing, but also additional WASP is recruited from the environment, highlighting a preference for the geometry found at the neck of 500 nm bead-induced invaginations (Fig. 3D). Finally, when analyzing the ventral surface of cells with 2D STED, we see a higher proportion of “U”-shaped invaginations with uniform WASP closer to the cell front and omega-shaped invaginations with punctate WASP further back (Fig. 3E), supporting our observations that over time the membrane reconfigures from open-necked to closed-neck and that WASP reorganizes as more favorable neck geometries arise (Fig. 3F).

### Dual inputs of membrane curvature and cell polarity inform WASP recruitment

WASP is consistently recruited to sites of inward membrane deformation. However, if WASP enriched everywhere the membrane was invaginated by the substrate, cells might engage with the coverslip across their entire surface and be unable to productively migrate. We next investigated whether there was spatial information restricting where WASP can respond to curvature.

Established curvature-sensitive proteins typically enrich to a given plasma membrane morphology irrespective of location in the cell (Zhao et al., 2017; Lou et al., 2019). Strikingly, WASP only enriched to beads in the front half of the cell (Fig. 4A). When a cell ran into a bead, WASP was recruited within seconds (Fig. 2B). Then, as the cell migrated over the bead and it progressed to the cell rear, WASP signal was lost (Fig. 4B-C; Fig. S5A). The disappearance of WASP correlated more strongly with puncta position than duration, as puncta that nucleated further from the cell front exhibited shorter lifetimes (Fig. S5B). To ensure that WASP disappearance was not a consequence of poor membrane deformation at the cell rear, we imaged the membrane and confirmed that 500 nm bead-induced membrane deformations persisted at the cell rear but lost WASP signal (Fig. 4D). Additionally, using 3D STED we observed that bead-induced invaginations lost WASP over time, as they moved toward the cell rear (Fig. S5C). Invaginations at the cell front were consistently WASP-positive, while invaginations at the rear of the same cells did not have WASP despite displaying similar membrane geometries (Fig. S5D). To our knowledge, this is the first report of a curvature-sensitive protein that integrates both membrane curvature and cell polarity.

**Fig. 4.**
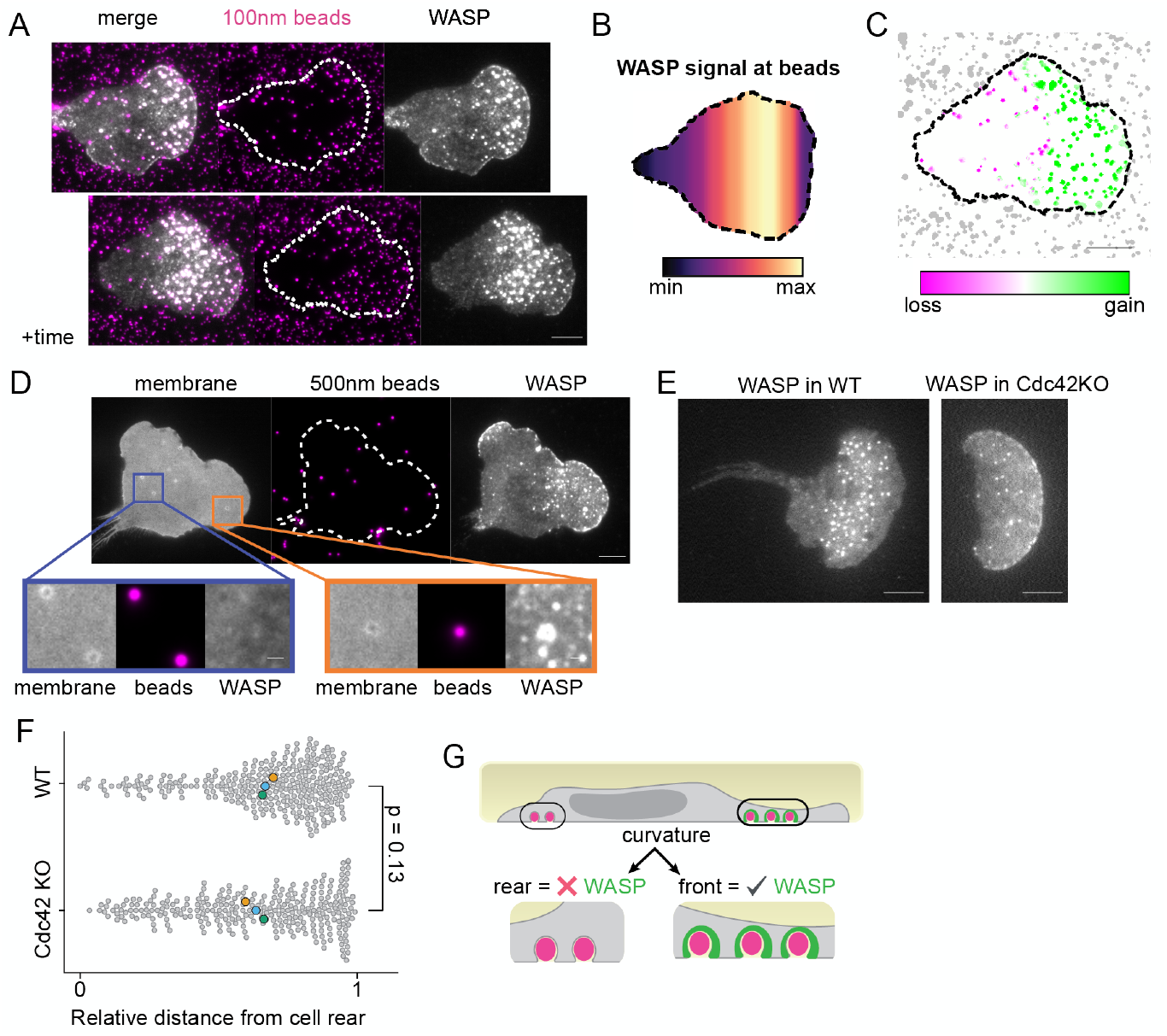
WASP recruitment depends on both curvature and cell polarity. A) WASP enriches to bead-induced membrane deformations only at the cell front. Beads that have signal in the top montage lose signal as they move towards the cell rear. Images are approximately 5 minutes apart. (B) Spatial distribution of average WASP signal from approximately 135 beads that traverse the cell length. Data was collected across three experiments. The average WASP signal at a bead peaks in the front 20 to 40% of the cell and then falls off despite continued presence of the bead under the cell. (C) Comparison of WASP signal gain and loss at beads from time point data in (A). Beads at the cell front gain WASP while beads at the cell rear lose WASP. (D) Membrane labeling confirms that beads continue to deform the membrane as they approach the cell rear yet no longer recruit WASP. Orange inset shows a bead at the front that deforms the membrane and recruits WASP, while the purple inset shows beads at the cell rear that deform the membrane but do not recruit WASP. Scale bar of the inset is 1 μm. (E) Exogenous WASP forms puncta in Cdc42 KO cells. (F) The position of WASP puncta appearance in both wild type and Cdc42 KO cells is biased toward the cell front and is not significantly different by an unpaired two-tailed t-test on replicate means: p = 0.13. More than 280 total puncta were measured for each cell background in 12 and 14 cells, respectively, that were collected across three experiments. Enlarged markers denote replicate means. Mean relative position of WASP puncta appearance is 0.68 ± 0.01 in wild type cells and 0.64 ± 0.02 in Cdc42 KO cells on the replicate level, with the cell rear defined as 0 and the cell front defined as 1. (G) Updated model to reflect that that membrane curvature alone is not sufficient to drive WASP recruitment. Scale bars are 5 μm unless otherwise specified.

To investigate how WASP combines cell polarity inputs with its curvature-sensitive membrane enrichment, we sought to better understand the inputs that specify the region permissive for WASP recruitment. WASP/N-WASP is known to link Cdc42 to Arp2/3 complex activation (Rohatgi et al., 1999; Higgs and Pollard, 2000), and Cdc42 activity is polarized towards the leading edge of migrating neutrophils (Yang et al., 2016). However, WASP puncta continued to form in a polarized fashion in Cdc42 knockout cells (Fig. 4E-F). Therefore, despite the strong link between Cdc42 and WASP, Cdc42 is not necessary for WASP foci formation or polarization in neutrophils. As an orthogonal approach to remove potential polarity input from both Cdc42GTP and RacGTP binding, we rescued WASP-null cells with a WASP mutant lacking the Cdc42- and Rac-interactive binding (CRIB) domain and observed similarly polarized puncta (Fig. S5E).

Our data show that WASP puncta formation and curvature sensitivity occur only in a permissive, front-polarized region. WASP identifies membrane invaginations as they form at the cell front and then leaves around the time they cross the cell midline (Fig. 4G). We next investigated how this polarization impacts the cytoskeleton.

### WASP links membrane topology with the cytoskeleton

To determine whether actin polymerization occurs at WASP puncta, we transduced WASP knock-in cells with a TagRFP-T-tagged Arp3 subunit of the Arp2/3 complex. Arp3 co-localizes with endogenous WASP puncta at the cell front (Fig. 5A-B). Unlike in CME where N-WASP signal peaks roughly ten seconds before that of Arp3 (Taylor et al., 2011), we were unable to resolve a temporal offset between recruitment of WASP and Arp3 at puncta. Additionally, in comparing the spatial distributions of WASP and Arp3 across the cell length, we find significant overlap between the signals (Fig. 5C). Given these observations, we conclude that Arp3 behavior closely mirrors that of WASP at puncta. Enrichment of an actin nucleator to these sites suggests polymerization is occurring and that native WASP puncta are filamentous actin (F-actin) rich structures.

**Fig. 5.**
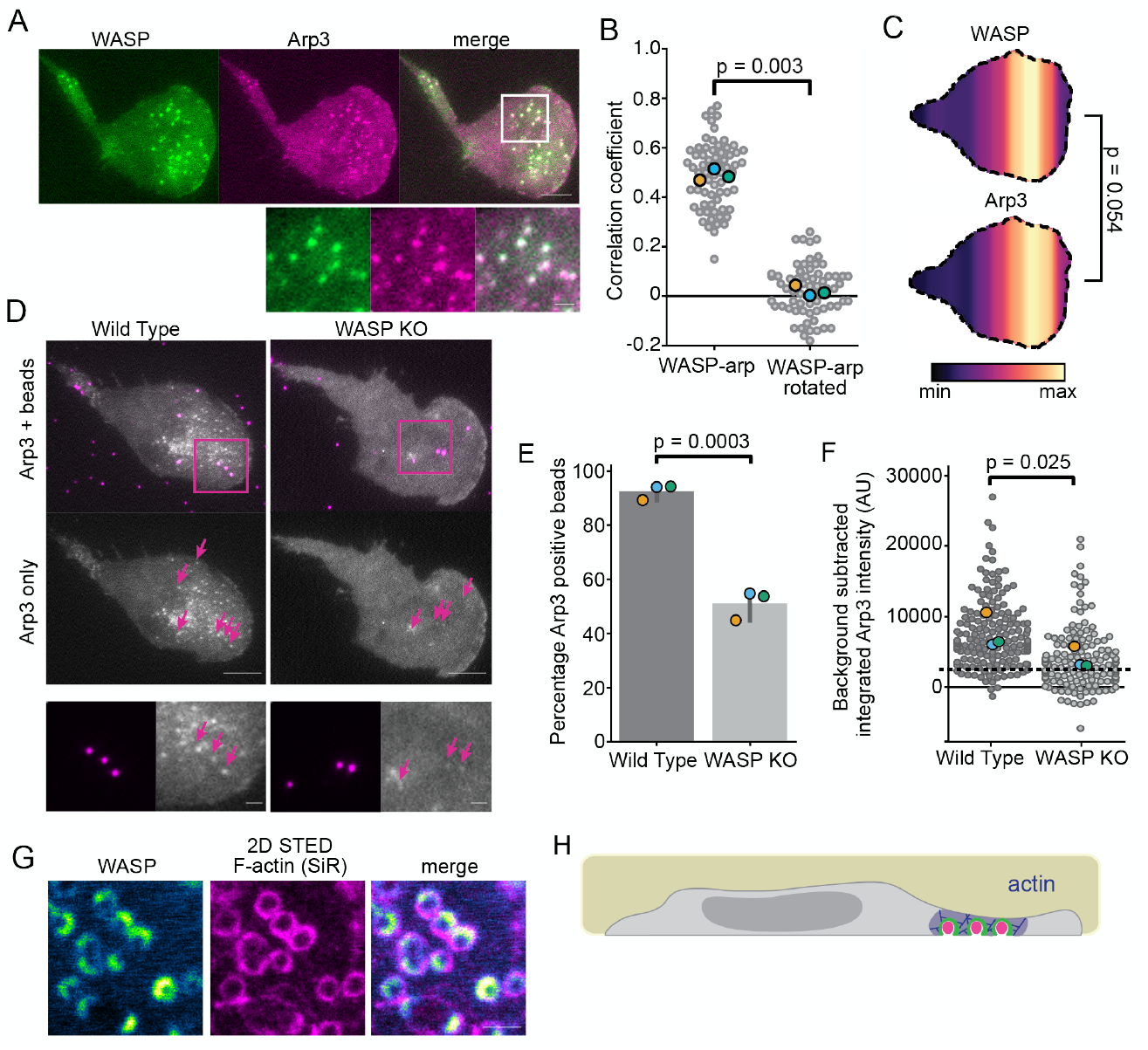
WASP mediates actin polymerization at sites of membrane deformation. (A) Endogenous WASP co-localizes with TagRFP-T-Arp3. Scale bars are 5 μm and 1 μm in the inset. (B) Pearson correlation coefficients of WASP and Arp3 show significant co-localization (r = 0.49 ± 0.01) compared to the correlation coefficients between WASP and a 90° rotation of the Arp3 channel (0.02 ± 0.02). n = 68 7.5X7.5 μm ROIs collected across three experiments. p = 0.003 by a paired two-tailed t-test on replicate means, which are denoted throughout the figure by enlarged markers. (C) Spatial distributions of WASP and Arp3 puncta reveal similar organization. n = 18 co-expressing cells collected across three experiments with n_WASP_ = 559 and n_Arp3_ =417 total puncta. The mean relative puncta position is 0.64 ± 0.02 for WASP and 0.65 ± 0.02 for Arp3, with the cell rear defined as 0 and the cell front defined as 1. p = 0.054 by a paired two-tailed t-test on the mean puncta position of each marker in each cell. (D) WASP-null cells fail to recruit appreciable amounts of the Arp2/3 complex to 100 nm bead-induced invaginations. Arrows denote the position of beads that are under the lamellipod. Below is a zoomed inset of the boxed region. All images are scaled equally. Scale bars are 5 μm and 1 μm in the inset. (E) Only about half of the beads (51.3 ± 3.1%) confined under lamellipodia in WASP KO cells are able to form Arp2/3 puncta (determined by the ability to fit a Gaussian to Arp3 signal at a bead) compared to 93.7 ± 1.7% of beads for wild type cells. n_WT_ = 172 beads and n_KO_ = 130 beads collected across three experiments. p = 0.0003 by an unpaired two-tailed t-test on replicate means. (F) Integrated intensity of background subtracted Arp3 signal at beads was calculated for the brightest Arp3 frame. The dashed line marks the approximate threshold needed to fit a Gaussian in panel (E). Measurements below this line are close to background or noise. The mean intensity of Arp3 at beads is 7696 ± 1454 AU in wild type cells and 4020 ± 886 AU in WASP KO cells on the replicate level. n_WT_ = 164 and n_KO_ = 143 beads collected across three experiments. p = 0.025 by a paired t-test on replicate means. (G) WASP and F-actin (reported with SiR actin) co-localize at bead-induced membrane invaginations at the cell front. Scale bar is 1 μm. (H) Updated model to reflect poIarized WASP-dependent actin polymerization.

Next, we sought to determine whether actin polymerization occurs at bead-induced invaginations, and, if so, whether this process depends on WASP. Confining our Arp3-tagged cells onto beads, we found that the Arp2/3 complex, like WASP, enriches to nearly all bead-induced invaginations at the cell front (Fig. 5D). However, repeating these measurements in WASP-null HL-60s led to a roughly 50 % reduction in the number of beads that induced Arp2/3 puncta formation and a significant decrease in the integrated Arp3 intensity at beads despite similar expression levels in both backgrounds (Fig. 5D-F). Additionally, while there persisted a band of Arp3 signal immediately behind the leading edge, Arp3 signal across the rest of the cell was quite uniform, and puncta outside of those formed at beads were rare (Fig. 5D). Therefore, the Arp2/3 complex is normally recruited to sites of inward membrane deformation through its association with WASP.

Finally, we used the live-cell actin probe SiR-actin to assay for F-actin at bead-induced invaginations and found co-localization of SiR-actin with WASP at beads under the cell front (Fig. 5G-H). Our finding that WASP plays an important role in linking membrane deformation to the actin cytoskeleton led us to next investigate how WASP-dependent actin structures affect neutrophil migration.

### Integration of topographical features during neutrophil migration depends on WASP

Given the clear *in vivo* phenotype of WASP-null neutrophils (Snapper et al., 2005; De Noronha et al., 2005; Westerberg et al., 2005; Jones et al., 2013) and our observation that WASP recruitment is regulated by inward membrane curvature, we hypothesized that WASP-dependent migration may be particularly acute for cells challenged by a complex, membrane-deforming environment. To asses this, we probed the role of WASP during migration on nanoridged substrates designed to mimic the aligned collagen fibers cells encounter in vivo (Ray et al., 2017).

First, we tested whether WASP puncta form at sites of inward curvature generated by nanoridges, as they did at beads (Fig. 2) and on collagen fibers (Fig. S2). Indeed, WASP puncta accumulate on nanoridges where the pattern pushes into the cell (Fig. 6A-B). Unlike focal adhesions that form on both the ridges and grooves of these patterns (Ray et al., 2017), WASP localized specifically to the positive (inward) curvature-inducing ridges. Additionally, despite long stretches of contact between cells and the patterns, WASP enrichment remained punctate and failed to continuously localize across ridges under the cell. The fact that curvature alone does not explain WASP patterning on nanoridges suggests that there may be other inputs determining the spatial scale of WASP self-organization (Li et al., 2012; Banjade and Rosen, 2014; Case et al., 2019).

**Fig. 6.**
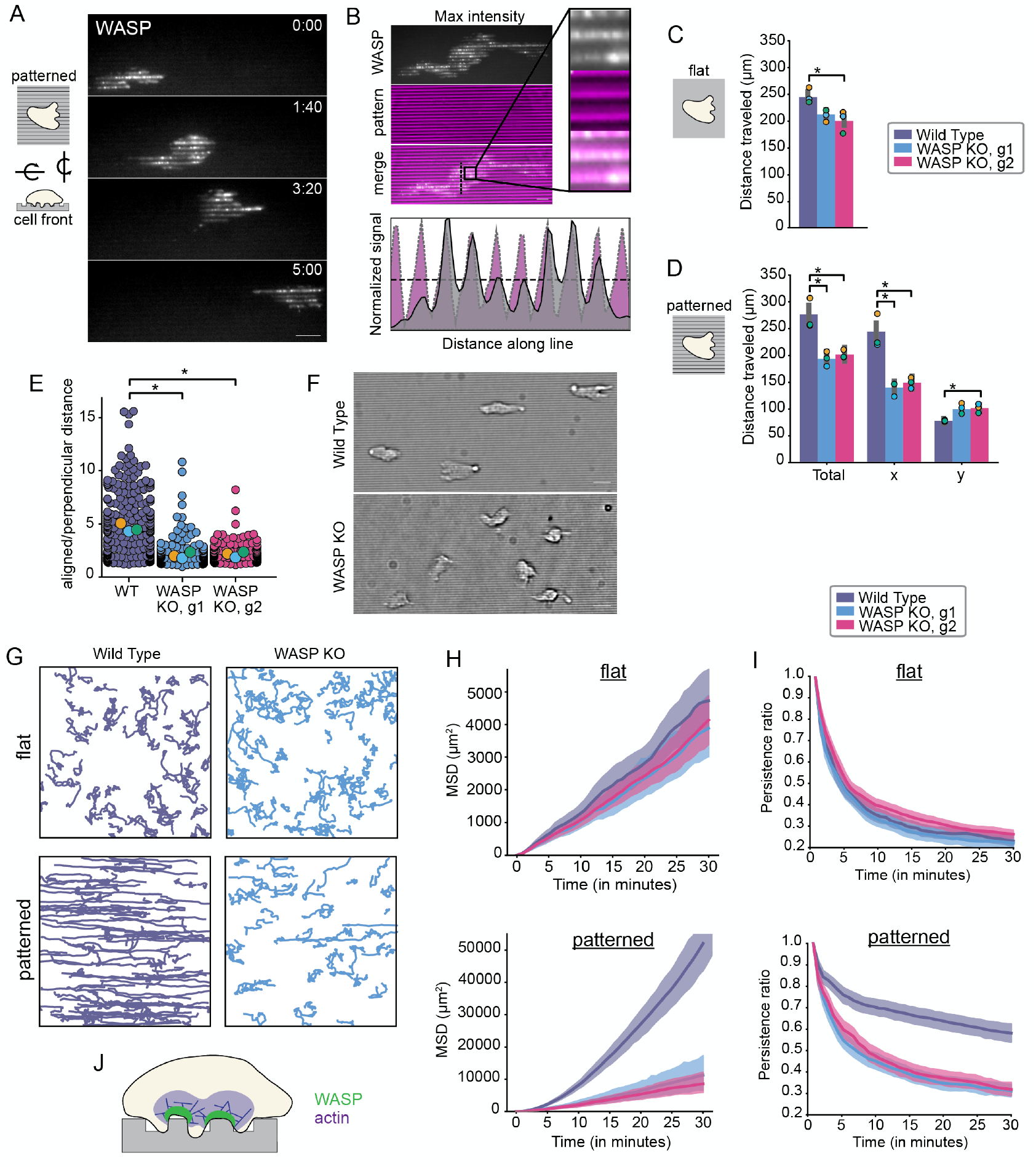
WASP-null cells are defective in their ability to interpret the topological features of their environment. (A) WASP puncta enrich at nanoridges as cells migrate over patterns. Scale bar is 10 μm. (B) Max intensity projection of data in (A) shows that WASP localizes to nanoridge peaks. An intensity profile for WASP and the pattern along the vertical line in the merged image is shown below. Scale bar is 5 μm and the inset is approximately 4×4 μm. (C) There is a modest difference in the total distance traveled by wild type (purple) and WASP KO (guide 1 = blue, guide 2 = magenta) cells plated on flat substrates. n_WT_ = 253 cells, n_KO1_ =215 cells, and n_KO2_ = 181 cells collected across three experiments. Total distance traveled: wild type = 246 ± 1 μm, WASP KO1 = 210 ± 6 μm, and WASP KO2 = 201 ± 12 μm on the replicate level. Replicate means are denoted throughout the figure by enlarged markers. Only one of the WASP KO clones was significantly different from wild type. For this case, p = 0.026 by a paired two-tailed t-test on replicate means. In the other case, p = 0.13. See also Video 6. (D) The total distance traveled by WASP KO cells is significantly less than that of wild type cells on nanoridged substrates due to defective movement along the patterns (here the x-axis). n_WT_ = 188 cells, n_KO1_ = 172 cells, and n_KO2_ = 242 cells collected across three experiments. Total distance traveled: wild type = 273 ± 17 μm, WASP KO1 = 194 ± 8 μm, and WASP KO2 = 202 ± 5 μm on the replicate level. Total distance traveled in x: wild type = 241 ± 19 μm, WASP KO1 = 139 ± 8 μm, and WASP KO2 = 149 ± 6 μm on the replicate level. Total distance traveled in y: wild type = 78 ± 1 μm, WASP KO1 = 102 ± 6 μm, and WASP KO2 = 102 ± 5 μm on the replicate level. Asterisk denotes p < 0.05, which was determined by a paired two-tailed t-test on replicate means. Unless labeled, the difference is not significant. p-values: p_total_,_WT/KO1_ = 0.018, p_total_,_WT/KO2_ = 0.028, p_x_,_WT/KO1_ = 0.023, p_x_,_WT/KO2_ = 0.023, p_y_,_WT/KO1_ = 0.061, p_y_,_WT/KO2_ = 0.035. See also Video 7. (E) Ratio of aligned to perpendicular migration reported in (D) highlights defective nanopattern sensing by WASP KO cell lines. Average ratio of total aligned to total perpendicular distance traveled: wild type = 4.02 ± 0.23, WASP KO1 = 1.47 ± 0.17, and WASP KO2 = 1.57 ± 0.15 on the replicate level. Asterisk denotes p < 0.05, which was determined by a paired two-tailed t-test on replicate means. p_WT/KO_ = 0.012 and p_WT/KO2_ = 0.008. (F) Bright field images reveal clear alignment of wild type cells to horizontal nanoridges while WASP-null cell largely fail to align. Scale bar is 5 μm. (G) Wild type and WASP-null cell trajectories over one hour for cells migrating on flat substrates (top) and nanoridged substrates (bottom). Image dimensions are approximately 665 × 665 μm. (H) The mean squared displacements (MSDs) of wild type and WASP-null cells are indistinguishable on flat substrates (top) while the MSDs of WASP-null cells are significantly less than that of wild type cells on nanoridged substrates (bottom). Shading denotes a 95% confidence interval. Trajectories are the same as those assayed in (C) and (D), respectively. Of note, the MSD of wild type cells is ten-fold greater on nanoridged substrates than on flat substrates. (I) The persistence ratios of wild type and WASP-null cells are indistinguishable on flat substrates (top) while the persistence ratios of WASP-null cells are significantly less than that of wild type cells on nanoridged substrates (bottom). Shading denotes a 95% confidence interval. Trajectories are the same as those assayed in (C) and (D), respectively. (J) Model of the role of WASP and WASP-mediated actin polymerization in topology sensing and contact guidance.

To determine whether WASP enrichment on nanoridges reflected a role in migration, we next compared the effect of flat versus textured substrates on the migration of wild type and WASP-null HL-60s (Fig. S1). There was a modest difference between the total distance travelled by wild type and WASP-null cells on flat substrates (15%-19% decrease) (Fig. 6C; Video 6). In contrast, WASP-null cells plated on nanoridges exhibited a greater reduction in total distance travelled compared to wild type cells (27-30% decrease) (Fig. 6D, Video 7). When we decomposed cell movement into axes that aligned with or were perpendicular to the nanopatterns, the observed migration defect was explained by a decrease in the ability of WASP-null cells to orient their migration along patterns. In fact, WASP-null cells moved 60% less than wild type cells in the direction of the patterns while moving 30% more than wild type cells in the direction perpendicular to the patterns. This is further reflected by a large difference in the ratio of aligned to perpendicular migration between wild type and WASP-null cells (Fig. 6E). The observed increase in perpendicular migration as opposed to a decrease in migration along both axes highlights a defect in the ability of WASP-null cells to sense and respond to the nanopatterned features of their substrate. Indeed, wild type HL-60 cells largely aligned with the ridges, while WASP-null cells did so with much lower frequency (Fig. 6F, Video 7). Wild type cells also moved more persistently along ridges and sometimes followed them across the entire field of view (>665 μm) (Fig. 6G, bottom left). In contrast, persistent migration along nanoridges occurred less frequently in WASP KO cells (Fig. 6G, bottom right).

To better understand how wild type and WASP-null cells differentially explore space on flat versus patterned substrates, we calculated the mean squared displacement (MSD) for each cell type in these different environments. In the absence of patterns, the MSDs of wild type and WASP KO cells were largely indistinguishable (Fig. 6H, top). However, on nanoridges the MSD was significantly larger for wild type cells than WASP KO cells across increasing time offsets (Fig. 6H, bottom). Due to an apparent difference in persistence between wild type and WASP KO cells on nanoridges (Fig. 6G, bottom; Video 7), we also calculated a persistence ratio, defined here as the ratio between displacement from the starting position and total distance travelled. In the absence of patterns the persistence ratio steadily falls off over increasing time windows for both wild type and WASP-null cells since there is no input biasing the direction of movement (Fig. 6I, top). Conversely, in the presence of nanoridges wild type cells are significantly more persistent than WASP-null cells on both short and long time scales (Fig. 6I, bottom).

Taken together, these observations reveal an essential role for WASP in substrate topology sensing and contact guidance in neutrophils (Fig. 6J).

## Discussion

Neutrophils need to navigate complex three-dimensional paths through dense extracellular matrices *in vivo* (Lämmermann et al., 2013). To move in this environment, a cell must be able to sense its physical surroundings and coordinate its movement with the features of its substrate. Our work suggests that WASP plays a role in mediating this process. First, WASP helps detect the topology of the substrate by enriching to sites of inward, substrate-induced membrane deformation (Fig. 2 and Fig. 3). WASP then facilitates recruitment of the Arp2/3 complex for local actin assembly, thereby coupling substrate features with the cytoskeleton (Fig. 5). WASP functions as a dual integrator of cell polarity and membrane curvature, enriching to membrane invaginations only at the cell front (Fig. 4). To our knowledge, this is the first report of curvature sensitivity that is informed by physical location within a cell. Through these characteristics, WASP is able to (1) identify substrate features on which to anchor the cell, (2) pattern actin polymerization to leverage substrate contact sites for locomotion, and (3) constrain substrate engagement to regions that yield productive migration. As a consequence of these features, WASP-null neutrophils are particularly defective at coordinating substrate topology with cell guidance (Fig. 6).

While the importance of WASP has long been appreciated due to its role in Wiskott-Aldrich syndrome, its contribution to neutrophil migration has been unclear. In animal models of this disease, neutrophils are defective at homing to sterile wounds (Jones et al., 2013) and sites of infection (Snapper et al., 2005; Kumar et al., 2012). However, in *vitro* WASP-null neutrophils have shown mixed phenotypes. Some studies have reported defective migration (Ochs et al., 1980; Fritz-Laylin et al., 2017) while others found no defect (Zicha et al., 1998; Zhang et al., 2006). One possible reason for this discrepancy is our finding that WASP is primarily used during migration in complex environments where substrate topology sensing and engagement play a critical role.

Much of what we know about WASP function has been inferred from that of its ubiquitously expressed homologue N-WASP, which plays a key role in clathrin-mediated endocytosis (Merrifield et al., 2004; Benesch et al., 2005; Kessels and Qualmann, 2002), among other functions (Isaac et al., 2010; Nusblat et al., 2011; Mizutani et al., 2002; Yamaguchi et al., 2005; Yu et al., 2012; Yu and Machesky, 2012; Zhang et al., 2006; Misra et al., 2007; Liu et al., 2013). Whether WASP plays a similar role in immune cells was not known. Our work reveals that a critical function of WASP in neutrophils is divergent from the canonical functions of N-WASP. However, we find some parallels between WASP and N-WASP; both proteins use membrane invagination to locally polymerize actin via Arp2/3 complex recruitment. This process culminates in clathrin-mediated endocytosis for N-WASP, but not WASP, where the function appears to be in coupling the actin cytoskeleton to substrate-induced membrane deformations that guide cell movement. This is not the first example of cells repurposing the curvature sensitivity of the endocytic machinery to help engage with their substrate. In particular, collagen fibers induce persistent non-endocytic clathrin-coated structures that aid in 3D migration in other cellular contexts (Elkhatib et al., 2017). We propose WASP is performing a similar function in highly motile neutrophils.

WASP and N-WASP have also been implicated in building podosomes and invadopodia (Isaac et al., 2010; Nusblat et al., 2011; Mizutani et al., 2002; Yamaguchi et al., 2005; Yu et al., 2012; Yu and Machesky, 2012), which are membrane exvaginations that participate in extracellular matrix degradation and mechanosensation (Albiges-Rizo et al., 2009; Murphy and Courtneidge, 2011). The WASP puncta we observe are triggered by membrane invagination, rather than exvagination, and wrap around the substrate, instead of poking into it. Therefore, WASP structures in neutrophils are distinct from invadopodia and podosomes and may represent a variation on how cells interact with their substrate. Also, rather than degrading ECM, neutrophils squeeze through pores in collagen matrices, which may rely on an alternate mode of substrate engagement (Lämmermann et al., 2013; Steadman et al., 1997; Allport et al., 2002; Lämmermann et al., 2008).

WASP is unusual in that it is able to integrate both cell polarity and membrane curvature for its recruitment; only membrane deformations in the cell front result in WASP enrichment (Fig. 4A-D, Fig. S5C-D). How does WASP sense cell polarity? An obvious candidate is Cdc42; it is one of the primary inputs to WASP-mediated actin assembly (Rohatgi et al., 1999; Higgs and Pollard, 2000), and Cdc42 activity is polarized towards the leading edge of migrating neutrophils (Yang et al., 2016). However, we find Cdc42 to be dispensable for WASP polarity (Fig. 4E-F). WASP can also bind Rac with a lower affinity (Kolluri et al., 1996), but structure-function analysis of WASP reveals that the Cdc42- and Rac-interactive binding domain of WASP is also dispensable for its polarity (Fig. S5E). Importantly, there are other inputs that activate WASP family proteins; N-WASP-mediated actin polymerization can be elicited in the absence of Cdc42 both in *vitro* (Takano et al., 2008) and by pathogens (Moreau et al., 2000). Therefore, future studies will be needed to elucidate the additional inputs to WASP’s polarity sensing. Similarly, the molecular basis of WASP’s curvature sensitivity remains unknown. Previous studies have suggested that the BAR domain-containing protein FBP17 plays a role in the curvature sensitivity of N-WASP (Lou et al., 2019; Tsujita et al., 2015), but we find that WASP can sense curvature in the absence of FBP17 in neutrophils (Fig. S3). Whether this is due to a difference between WASP and N-WASP or whether the use of dominant-negative FBP17 in initial studies inhibited other N-WASP-associating curvature-sensitive proteins, which can normally co-oligomerize and compensate for one another (Chan Wah Hak et al., 2018; Tsujita et al., 2013), is unknown.

Overall, our work establishes WASP as a link between cell movement and substrate topology. Previous reports have found that different cell types, including HL-60s, can sense nanopatterned topologies and respond to them through reorientation and aligned migration (Driscoll et al., 2014; Sun et al., 2015). In the case of adherent cells, this is thought to be through imposing spatial constraints on focal adhesion formation and maturation (Ray et al., 2017). However, for loosely adherent cells like neutrophils and *Dictyostelium discoideum* that do not rely on long-lived adhesive structures (Lämmermann and Sixt, 2009; Driscoll et al., 2014), the molecular mechanism unpinning physical environment sensing and alignment to substrate topologies has remained unclear. Some proposed mechanisms for cytoskeletal remodeling in response to membrane deformation include membrane geometry sensing (Itoh et al., 2005; Zhao et al., 2017; Lou et al., 2019; Takenawa and Suetsugu, 2007), membrane tension sensing (Houk et al., 2012; Tsujita et al., 2015; Diz-Muñoz et al., 2016), force adaptation (Bieling et al., 2016; Mueller et al., 2017), and even actin filament bending (Risca et al., 2012). The fact that WASP is both curvature-sensitive and essential for HL-60 alignment to nanoridges supports WASP-mediated actin polymerization as a mechanism by which loosely adherent cells can sense and align with their substrate’s topology.

Membrane curvature has emerged as a key regulator for WASP family proteins; other NPFs from this family such as N-WASP and WAVE also preferentially enrich to sites of membrane curvature (Lou et al., 2019; Pipathsouk et al., 2019). Unique curvature preferences among WASP family NPFs could help differentiate their organization and inform the morphology of the actin networks they generate. For N-WASP, enrichment to membrane invaginations could facilitate scission of endocytic vesicles from the plasma membrane (Kaksonen et al., 2005; Akamatsu et al., 2020). For WAVE, saddle preference could help organize a coherently advancing flat lamellipod (Pipathsouk et al., 2019). And for WASP, enrichment to stable inward curvature (in particular the saddle-shaped necks of invaginations) could help the cell integrate substrate topologies with motility. Interestingly, these proteins also appear to have different rules for reading curvature—for instance WASP is much better than WAVE at persistently enriching to saddles at bead-induced membrane invaginations (Fig. 2F-G). Whether these individualized responses arise from subtle differences in saddle-curvature preference, WAVE complex’s extinction at non-motile barriers (Weiner et al., 2007), or WASP’s broader permissive zone (Fig. 1A) will be an interesting avenue for future investigation. Notably, in some contexts WASP can even substitute for WAVE (Veltman et al., 2012; Zhu et al., 2016; Tang et al., 2013), suggesting overlap in the membrane geometries they can read. Future studies systematically investigating the curvature preferences among WASP family NPFs will expand our understanding of how variation in membrane geometry can feed into the diversification of actin network patterning.

## Materials and methods

### Cell Culture

HL-60 cells are from the lab of Henry Bourne. PLB-985 cells were originally obtained from Onyx Pharmaceuticals. RNA-seq was recently performed on both of these backgrounds (Rincón et al., 2018), confirming cell line identity and supporting a previous report that PLB-985 cells are actually a sub-line of HL-60 cells (Drexler et al., 2003). Both lines were grown in RPMI 1640 media supplemented with L-glutamine and 25 mM HEPES (Corning, MT10041CV) and containing 10% (v/v) heat-inactivated fetal bovine serum (Gibco, 16140071) and 1X penicillin-streptomycin (Gibco, 15140148) (called HL-60 media). Cultures were split to 0.17-0.2 million cells/mL every two to three days and grown at 37°C/5% CO2. Cells were differentiated for experiments by diluting cells to 0.2 million/mL, adding 1.5% (v/v) DMSO (Santa Cruz Biotechnology, 358801), and incubating for 4-5 days. HEK293T cells (used to generate lentivirus for transduction of HL-60s) were grown in DMEM (Gibco, 11995065) containing 10% (v/v) heat-inactivated fetal bovine serum (Gibco, 16140071) and 1X penicillin-streptomycin (Gibco, 15140148) and maintained at 37°C/5% CO2. Cells were routinely checked for mycoplasma contamination.

### Plasmids

A vector for mammalian expression of TagRFP-T-Arp3 was generated by PCR amplification of the Arp3 coding sequence and Gibson assembly into a lentiviral pHR backbone containing a N-terminal TagRFP-T sequence. The sequence for mNeonGreen2_1___10_ (Feng et al., 2017) was similarly transferred to a pHR backbone. Hem1-eGFP, used for measuring WAVE complex recruitment, was previously described (Diz-Muñoz et al., 2016; Pipathsouk et al., 2019) and similarly generated by transferring the protein coding sequence to a pHR backbone.

For generating knockout lines, guide RNAs with homology to exon 1 of *WAS* (5’ CCAATGGGAGGAAGGCCCGG 3’ and 5’ GCTGAACCGCTGGTGCTCCT 3’) and exon 4 of *FNBP1* (5’ ACGAAATGAATGATTACGCA 3’) were selected using the CRISPR design tool in Benchling (www.benchling.com) and cloned into the previously described LentiGuide-Puro vector (Addgene plasmid 52963) (Sanjana et al., 2014). The vector used to express human-codon-optimized *Streptococcus pyrogenes* Cas9-BFP was also previously described (Graziano et al., 2017).

For endogenous tagging of WASP with plasmid donor, 500 base pairs of the 5’ untranslated region and the start codon (5’ homology arm), TagRFP-T flanked by unique seven amino acid long linkers, and the 500 base pairs following the start codon (3’ homology arm) were Gibson assembled into a minimal backbone (pUC19). A similar approach was used for endogenous tagging of clathrin light chain A *(CLTA).*

### Transduction of HL-60 cells

HEK293T cells (ATCC) were seeded into 6-well plates and grown until approximately 70% confluent. For each well, 1.5 μg pHR vector (containing the appropriate transgene), 0.167 μg vesicular stomatitis virus-G vector, and 1.2 μg cytomegalovirus 8.91 vector were mixed and prepared for transfection using TransIT-293 Transfection Reagent (Mirus Bio, MIR 2705). Following transfection, cells were incubated for 3 days, after which virus-containing supernatants were harvested and concentrated approximately 40-fold using Lenti-X Concentrator (Takara Bio, 631232) per the manufacturer’s instructions. Concentrated viruses were frozen and stored at 80°C until needed or used immediately. For all transductions, virus was mixed with 0.36 million cells in growth media supplemented with 8 μg/mL Polybrene (Sigma-Aldrich, H9268) and incubated overnight. Fluorescence-activated cell sorting (FACS) (BD Biosciences, FACSAria2 or FACSAria3) was used to isolated cells with the desired expression level of the transgene.

### Generation of CRISPR knockout HL-60 cell lines

WASP and FBP17 HL-60 knockout cell lines were generated as described in (Graziano et al., 2019). Briefly, HL-60s were transduced with a puromycin-selectable vector containing an sgRNA sequence for the gene of interest. Following puromycin selection, cells were transduced with a *S. pyrogenes* Cas9 sequence fused to tagBFP. Cells expressing high levels of Cas9-tagBFP were isolated with FACS. Cells were then diluted into 96-well plates at a density of approximately one cell per well in 50% (v/v) filtered conditioned media from a healthy culture, 40% (v/v) fresh HL-60 media, and 10% (v/v) additional heat-inactivated fetal bovine serum. Clonal cell lines were expanded and validated using amplicon sequencing and immunoblot. Two clonal lines were generated for WASP from separate CRISPR assemblies (guide 1: 5’ CCAATGGGAGGAAGGCCCGG 3’ and guide 2: 5’ GCTGAACCGCTGGTGCTCCT 3’) and one clonal line was generated for FBP17 (5’ ACGAAATGAATGATTACGCA 3’).

The Cdc42 PLB-895 knockout line was generated as described in (Bell et al., 2021). Briefly, PLB-895 cells were electroporated with a Cas9-sgRNA ribonucleoprotein (RNP) complex. Tandem guides targeting exon 4 of Cdc42 were purchased from Synthego (5’ TTTCTTTTTTCTAGGGCAAG 3’ and 5’ ATTTGAAAACGTGAAAGAAA 3’). Guides were complexed with Cas9 protein and electroporated into cells using a custom suspended-drop electroporation device (Guignet and Meyer, 2008).

### Immunoblot assays

Protein content from one million cells was extracted using TCA precipitation. Samples were separated via SDS-PAGE and transferred to a nitrocellulose membrane. Membranes were blocked for approximately one hour in either a 1:1 solution of TBS (20 mM Tris, 500 mM NaCl [pH 7.4]) and Odyssey Blocking Buffer (LI-COR, 927-70001) or PBS with 1% BSA. For detection of WASP, the membrane was incubated overnight at 4°C in primary antibody (Cell Signaling Technology, 4860) diluted 1:1,000 in a 1:1 solution of TBST (TBS + 0.2% w/v Tween 20) and Odyssey Blocking Buffer. For detection of FBP17, the membrane was incubated for one hour at room temperature in primary antibody (a gift from P. De Camilli, Yale University) diluted 1:5,000 in PBST with 1% BSA. Membranes were washed three times with either TBST or PBST and then incubated for 45 minutes at room temperature with LI-COR secondary antibodies diluted 1:20,000 in either Odyssey Blocking Buffer or PBS with 1% BSA. Membranes were then washed three times with TBST or PBST, washed a final time with TBS or PBS, and imaged using an Odyssey Fc (LI-COR).

GAPDH (Invitrogen, MA5-15738) was used as a loading control for each blot and subjected to the same conditions as the antibody detecting the protein of interest.

### Generation of CRISPR knock-in HL-60 cell lines

Wild type HL-60 cells were transduced with a plasmid containing the non-fluorescent 1-10 segment of mNeonGreen2 (mNG2_1___10_) (Feng et al., 2017). Cells expressing the transgene were identified through electroporation (Lonza, 4D-Nucleofector) with a mCherry-conjugated mNeonGreen2_11_ (mNG2_11_) mRNA. Positive cells (those that were fluorescent in both 488 and 561) were identified using FACS (Sony, SH800) and a subset of cells that appeared to have a single insertion of mNG2_1___10_ was isolated. After a few days the mRNA was degraded and the cells lost fluorescence. This created the base mNG2_1___10_ HL-60 cell line used with mNG2_11_ knock-in.

sgRNAs of the target gene were obtained by *in vitro* transcription as described in (Leonetti et al., 2016) with modifications only to the PCR polymerase used (Phusion; New England Biolabs, M0530S). A 100 μL reaction of mNG2_1___10_ HL-60 cells were electroporated (Invitrogen, Neon) with a Cas9-sgRNA complex and a single-stranded DNA ultramer (ssDNA) (Integrated DNA Technologies) donor containing mNG2_11_, a linker, and 55 base pairs of homology to each side of the cut site. Cas9-sgRNA complex formation and electroporation protocols were adapted from (Brunetti et al., 2018; Garner et al., 2020). Briefly, 90 pmol of Cas9 containing a nuclear localization signal (QB3 MacroLab) and 270 pmol of sgRNA were mixed on ice and then incubated at room temperature for 30 minutes. Meanwhile, 2 million cells were spun down at 300xg for 5 minutes and washed with phosphate buffered saline to remove extracellular proteins. When the Cas9 complex was ready, cells were again spun down and this time resuspended in 100 μL room temperature R Buffer (Invitrogen, MPK10096). 100 pmol of ssDNA donor was added to the Cas9-sgRNA complex and the whole mixture was added to the resuspended cells. Samples were then electroporated at 1350V for 35 ms and recovered in 5 mL warmed media. Cells were monitored and allowed to recover. Using 100 μL tips led to recovery within only a few days. Cells were then expanded for selection with FACS. The percent of fluorescent cells ranged from 0.1-1%. For WAS, which has only one allele, the polyclonal line resulting from FACS sorting was sequence-verified and used for experiments. For other knock-ins, cells underwent clonal dilution as outlined above and surviving clones were screened using a PCR-based gel shift assay. Samples showing biallelic insertion were gel extracted and Sanger sequenced.

In the case of full-length fluorophore knock-in, plasmid donor was used in place of ssDNA donor. The above protocol was executed with only slight changes. First, 10 μL tips were used in place of 100 μL tips to remove the need for large amounts of donor. RNP components and the number of cells electroporated were therefore scaled down by a factor of ten as originally done in (Brunetti et al., 2018; Garner et al., 2020). Next, 800 ng of plasmid donor was provided in place of the ssDNA donor. Finally, post electroporation cells were rescued in 500 μL pre-warmed media in a 24 well plate and allowed to recover for ten to fourteen days. Despite lowered viability and longer recovery times post electroporation compared to 100 μL tips, similar taggingefficiencies were observed with this strategy.

N-terminal tagging was performed for WASP (WAS), clathrin LCa *(CLTA)*, and FBP17 *(FNBP1).* We used the following guide sequences to target near the start codon: *WAS* - 5’ GGCAGAAAGCACCATGAGTG 3’, *CLTA* - 5’ GAACGGATCCAGCTCAGCCA 3’, *FNBP1* 5’ - CGTCCCCTGCACCATGAGCT 3’.

### Compression of HL-60s for imaging

For all experiments excepting the STED, collagen, nanopatterns, and EDTA treatment, HL-60s were confined based on the method described in (Bell et al., 2018). For this work, a solution of 2% low-melt agarose (Gold Biotechnology, A-204) was made in L-15 media (Gibco, 21083-027) and microwaved in a loosely capped conical placed in a water reservoir. Heating was done in short increments to promote melting while preventing the solution from boiling over. Once completely melted, the gel was kept at 37°C to allow cooling while preventing solidification. Meanwhile, 500 μL of differentiated HL-60 cells were spun down at 200 × g for 5 minutes and then concentrated three-fold in HL-60 media. 5 μL of the concentrated cells were placed in the center of a circular well (Greiner Bio-One, 655891) and allowed to settle. When circular-welled plates could not be used, a 5 mm circular mold was inserted into the well. After the agarose had cooled to 37°C, 195 μL of agarose was slowly pipetted into the bottom edge of the well, allowing the agarose to spread over the droplet of cells. The pipette was then slowly raised up the side of the well to continue depositing agarose without sweeping away the covered cells. The plate was then allowed to dry uncovered for 20-45 minutes in an oven set to 37°C without humidity to promote high compression of the cells as in (Garner et al.,. Adequate flattening of the lamellipod was determined by observation of the complete cell footprint in TIRF.

Compression was also used in STED experiments, but modifications were used to accommodate imaging chambers (Ibidi, 80827). Briefly, a 1% solution of liquid agarose (Thermo Scientific, 17850) was made and poured into 8X8X5 mm square molds. After solidification, agarose pads were stored in L-15 media (Gibco, 21083-027) at 4°C. Cells were compressed by gently placing a room temperature agarose pad atop the well with tweezers and putting the chamber lid on top of the pad. Flattening was confirmed by cellular morphology in the confocal imaging mode.

The EDTA experiments required inelastic confinement to maintain the polarity and migration of cells. To achieve this, we confined cells between two glass surfaces as descried in (Malawista et al., 2000; Graziano et al., 2017). Briefly, glass slides and coverslips were thoroughly cleaned with Sparkleen laboratory detergent (Fisherbrand, 04-320-4) and then sonicated for 10 minutes first in acetone, then twice in ethanol, and finally three times in MilliQ water. The glass was immediately dried using N2 gas and stored in a clean, dry place. For experiments, differentiated HL-60s were concentrated to 20 million cells/mL in HL-60 media and a final concentration of 10 mM EDTA (Teknova, E0306) was added to “+EDTA” samples. 2 μL of cells were then placed onto a clean slide and a clean 18×18 mm coverslip was dropped on top. We confirmed that the cell solution was wicked across the majority of the coverslip surface, ensuring adequate confinement. Chambers were then sealed with VALAP (a mixture of equal parts Vaseline, paraffin wax, and lanolin) (Cold Spring Harbor Laboratory, 2015) and imaged.

### Cell compression onto beads

Agarose-based compression was done as described above with the addition of red (580/605) fluorescent carboxylate-modified microspheres (Invitrogen, F8887) to concentrated cells prior to plating. For quantitation of WASP and Arp2/3 recruitment to beads, a low density of beads was used (1:10,000 to 1:100,000). For bead carpets, as shown in Fig. 4A, beads were diluted only 1:1000. Since bead density is proportional to the bead volume a less stringent dilution was used for 500 nm beads (1:100).

### STED microscopy of cells on beads

All STED microscopy experiments were performed in plain glass 8-well chambers (Ibidi, 80827). Wells were coated with 100 μL of 20 μg/mL porcine fibronectin (prepared from whole blood) diluted in PBS at 37°C for 30 min and washed 3x using L-15 media (Gibco, 21083-027). Next, any residual imaging media was removed and 100 μL of beads diluted in L-15 were added to the well. We used either red-fluorescent (580/605) carboxylate modified beads of 200 or 500 nm diameter (Invitrogen, F8887) or non-fluorescent carboxylate modified beads of 500 nm diameter (Invitrogen, C37481). Bead density and adhesion were checked on the microscope prior to the addition of cells. Next, 800 μL of differentiated HL-60 cells were spun down at 200 × g for 5 minutes and resuspended in 100 μL of 5x concentrated CellMask Deep Red (Invitrogen, C10046) in L-15. Immediately after resuspension in the labeling solution, cells were spun down again at 200 × g for 5 minutes. After complete removal of supernatant, cells were resuspended in 100 μL of L-15 media and added to the well containing beads. To promote cell adhesion, the sample was placed in a 37°C incubator for 15 min. Finally, an agarose pad was placed atop cells for confinement and samples were imaged at room temperature.

### Cell migration on nanopatterned substrates

Nanopatterned and flat control substrates were purchased in 96-well plate format from Curi Bio (ANFS-0096). Wells were coated with 50 μL of 10 μg/mL porcine fibronectin (prepared from whole blood) dissolved in PBS for one hour at room temperature. The fibronectin solution was then removed and replaced with differentiated HL-60s (wild type or WASP KO) diluted by a factor of five into fresh HL-60 media containing one drop of the nuclear dye NucBlue (R37605, Thermo Fisher) per 500 μL. The plate was then transferred to a 37°C/5% CO2 incubator for 15 minutes to allow cells to adhere. The media was then gently pipetted over cells a few times to remove unattached cells and the media was exchanged for 100 μL fresh HL-60 media without NucBlue. The plate was then transferred to the microscope, which had been pre-heated to 37°C and an environmental chamber supplying 5% CO2 was turned on. The plate was allowed to equilibrate for one hour prior to imaging, giving cells time to align on nanopatterns. Wild type and WASP KO cells were always plated in adjacent wells, so that three positions could be taken per well over the time interval between frames (45s). Acquisition lasted one hour, and both bright field and 405 (nuclei) were collected.

### Cell migration in fluorescent collagen matrices

30 μL of pH-adjusted FITC-conjugated bovine skin collagen at 1 mg/mL (Sigma-Aldrich, C4361) was polymerized in a round, glass-bottomed well (Greiner Bio-One, 655891) for 3 minutes at room temperature and then gently washed with PBS (Elkhatib et al., 2017). 100 μL of differentiated HL-60s in HL-60 media were immediately added and allowed to migrate into the network up to one hour before imaging. WASP co-localization with fibers was assessed using a cell line that had WASP endogenously tagged with TagRFP-T.

### Microscopy

Total internal reflection fluorescence (TIRF) imaging experiments in Fig. 1 (excepting F), Fig. 2, and Fig. 4 (excepting E) were performed at room temperature using the ring TIRF light path of a DeltaVision OMX SR microscope (GE Healthcare) with a 60x/1.42 numerical aperture (NA) oil Plan Apo objective (Olympus) and 1.518 refractive index oil (Cargille). The system is equipped with 405, 445, 488, 514, 568, and 642 nm laser lines. mNeonGreen2 and eGFP-tagged proteins were imaged with the 488 nm laser line and Tag-RFP-T-tagged proteins were measured with the 568 nm laser line. The system was controlled with OMX AquireSR software (GE) and alignment of two channel images was performed with SoftWoRx (GE).

TIRF and confocal imaging experiments in Fig. 1F, Fig. 4E, Fig. 5 (excepting G), and Fig. 6 were performed at 37°C on a Nikon Eclipse Ti inverted microscope equipped with a motorized laser TIRF illumination unit, a Borealis beam conditioning unit (Andor), a CSU-W1 Yokugawa spinning disk (Andor), a 60X PlanApo TIRF 1.49 numerical aperture (NA) objective (Nikon), an iXon Ultra EMCCD camera (Andor), and a laser module (Vortran Laser Technology) equipped with 405, 488, 561, and 642 nm laser lines. NucBlue was measured with the 405 nm line, mNeonGreen2-tagged proteins were measured with the 488 nm line, and TagRFP-T-tagged proteins were measured with the 561 nm line. The system was controlled with μManager software (version 2.0-beta) (Edelstein et al., 2010).

Confocal microscopy and stimulated emission-depletion (STED) microscopy for Fig. 3 and panel 5G were performed at room temperature using an Abberior Expert Line system (Abberior Instruments GmbH) with an inverted Olympus IX83 microscope, QUADScan Beam Scanner (Abberior Instruments GmbH) and an Olympus UPlanSApo x100/1.41 oil immersion objective lens. The system was controlled with ImSpector (Abberior). Confocal measurements of mNeonGreen2 were made using a 488 nm line. All STED measurements were made using a 640 nm line and a 775 nm STED depletion laser. Sequential imaging was used to avoid cross-excitation. 2D STED imaging of the CellMask Deep Red and SiR-actin were performed using a 30 nm pixel size for X and Y. 3D STED imaging of CellMask Deep Red was performed in XZY mode using a z piezo stage (P-736 Pinano, Physik Instrumente), the Abberior adaptive illumination module RESCue, and a 30 nm voxel size for X, Y and Z (Staudt et al., 2011). The STED and confocal channels were aligned using measurements of 100 nm fluorescent beads on the Abberior auto-alignment sample. Spatial resolution of 3D STED was determined by imaging 40 nm far-red fluorescent beads purchased from Abberior.

### Image Analysis

Images were displayed in FIJI and WASP puncta and nuclei tracking were performed using the plug-in TrackMate (Tinevez et al., 2017). Tracks were filtered for desired property (duration, quality, etc.) within TrackMate and coordinates were exported as csv files and analyzed with custom Python code that made heavy use of the scikit-image package (van der Walt et al., 2014). All analysis code and a csv of all data are available at https://github.com/rachel-bot/WASP. Kymographs were created using the plug-in KymographBuilder (Mary et al., 2016).

### Quantification of WASP distribution

A single image from the indicated number of cells over three experiments was rotated using an interactive rigid transformation in FIJI so that cells were consistently oriented from left to right. Otsu thresholding was used to create a binary cell mask. The cell was then split into four regions: the leading edge, cell front, cell rear, and uropod. The leading edge was defined as the area removed by erosion of the front 40% of the cell mask with a 20×20 structuring element, which was determined to robustly isolate signal at the leading edge. The cell front was defined as the front 50% of the cell excluding the leading edge. The cell rear was defined as the region behind the front 50% of the cell and before the rear 20% of the cell, which was defined as the uropod. WASP signal in each region was determined by applying an Otsu threshold to a Difference of Gaussians (DoG) image (σ_high_ = 2, σ_low_ = 1), which highlights puncta. The signal in each region was measured by summing the binary of the thresholded DoG image. These values were then normalized to the total number of pixels in the cell mask for each region. Conversely, calculating the normalized integrated intensity in each region, after applying an Otsu threshold to remove background signal, yielded similar results.

### Spatial histogram analysis

Using cells oriented from left to right as explained above, a cell mask and its contour were created. The left, right, top and bottom bounds of the cell outline were extracted. The width of the cell in x was divided into the specified number of bins, here ten. Next, a DoG image of WASP or clathrin or Arp3 was generated and an Otsu threshold was applied. The user confirmed that this scheme of puncta identification was sufficiently robust. The resulting binary image was indexed and puncta centroids were recorded. The x-coordinate of each puncta was then used to assign it to one of the bins made from the cell mask’s x coordinate range. Similar analysis was repeated over all cells. Puncta counts per bin (each with different x-coordinates but representing the same portion of the cell, i.e. the front 1/10th) from all cells were then combined and normalized to the total puncta counts to get a density. For display purposes, this 1D array was extended in y to be 2D and multiplied by a binary mask of a well-polarized cell. All values outside of the cell body were converted to NaN to give a white background, and the cell contour was plotted on top to denote this boundary.

Similar analysis was employed to investigate the disappearance of WASP puncta in +/−EDTA conditions and the appearance of WASP puncta in wild type and Cdc42 knockout cells. To do this, puncta tracks were determined in FIJI using TrackMate (Tinevez et al., 2017), discarding traces that did not disappear during the course of the movie for the EDTA experiments or traces that were present at the start of the movie for the Cdc42 experiment. All tracks were manually checked to ensure they accurately captured the last or first frame of the puncta, respectively. Centroids at these times points were then passed to the analysis described above and their position relative to the cell length was determined by manual specification of the cell front and cell back in each frame using the matplotlib (Hunter, 2007) command “ginput” to select these landmarks.

### Quantification of WASP and Arp3 signal at beads

Single frames from movies of cells migrating over beads were selected based on the presence of beads under the lamellipod. The matplotlib (Hunter, 2007) command “ginput” was used to manually select beads that were completely under the lamellipod. The immediate region around the bead was isolated and a 2D Gaussian was fit to the diffraction-limited WASP or Arp3 signal. Intensity of the signal at the bead was calculated using the formula for volume under a Gaussian: 2πAσ_x_σ_y_, where the amplitude (A) and the standard deviation in each direction (σ_x_, σ_y_) come from the fit. This quantity is independent of background signal. For comparing WASP intensity across bead sizes, Gaussian volumes were normalized to bead surface area in accordance with our observation from STED microscopy that WASP signal is distributed across the entire invagination surface at sufficiently high curvatures (≤ 1/200 nm). For assessing Arp3 recruitment to beads in wild type and WASP KO cells, Gaussian volumes were not directly compared since many beads failed to induce Arp3 puncta in the absence of WASP and therefore could not be fit. Instead, we reported the fraction of beads able to form Arp3 puncta for each cell background. To avoid losing the intensity data entirely, we took an orthogonal approach and calculated a background subtracted integrated intensity for Arp3 signal at beads. To do this, Arp3 signal was tracked over time, and the frame with the brightest Arp3 signal in the region of the bead of interest was isolated. Arp3 signal was then masked with the bead signal to separate out the foreground and background. The mean background signal was calculated and subtracted from each pixel in the masked image. The resulting values were then summed to get the background subtracted Arp3 integrated intensity.

### WASP signal as a function of distance from the cell front

Movies where a cell moved the majority of its length over a bead without dislodging it from the TIRF plane were analyzed. At each time point, the cell rear and cell front were manually selected with the matplotlib (Hunter, 2007) command “ginput” to find the cell’s major axis. Beads that passed under the cell were selected and, at each time point, their (stationary) position was projected onto the major axis. This value was normalized to the major axis length in each frame to get a relative distance from the cell front (value of 0 to 1). For each time point, WASP intensity was estimated by summing the signal in a small box around the bead. The exactness of this measurement is not imperative since all traces are normalized to their maximum WASP signal to build a profile of relative signal as a function of distance from the cell front.

To get a population level profile, relative distances were discretized to the nearest 0.1 and traces from many beads were averaged to create a line plot. To better represent the data underlying this line, WASP data points in each positional bin were overlaid onto the plot and colored by the kernel density estimate of each value compared to all WASP values in the same positional bin.

### Co-localization by correlation analysis

Co-localization of curvature sensors with beads and WASP with clathrin/Arp3/FBP17 were determined by calculating the Pearson correlation coefficient between the flattened vectors of each channel. For statistical comparison of enrichment between curvature sensors, distributions of correlation coefficients were compared using an unpaired two-tailed t-test. For statistical comparison of co-localization between markers in the same cell, a negative control was generated by rotating one channel 90 degrees to remove co-localization while maintaining any punctate organization in the image (Dunn et al., 2011). The Pearson correlation coefficient was again calculated. The values of non-transformed and transformed correlation coefficients were combined across cells and experiments to create distributions. The means from each technical replicate were then compared between these two conditions using a paired t-test.

### Cell tracking and measurement of migration properties

Cell nuclei were tracked in FIJI using TrackMate (Tinevez et al., 2017) and resulting csv files were exported for analysis in Python.

To prevent double counting of cells, only traces that lasted at least half of the movie were kept. Then, to make sure all cells were compared over the same time window, only cells present for the first half of the movie were analyzed. Total distance traveled by each cell was calculated by applying the distance formula to each time point and summing the resulting displacements. This was performed for either both x and y or x or y independently, depending on the kind of distance being reported.

Mean squared displacement (MSD) was calculated using 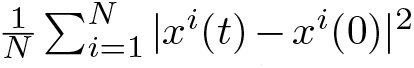 where *x^i^*(*t*) is the position of *i*-th cell at time *t* and *x*^(*i*)^(0) is its initial position. *N* denotes the total number of cells.

The persistence ratio for each cell was calculated using 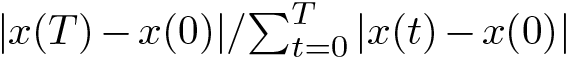 where *T* denotes the time point being compared with *t* = 0 (Gorelik and Gautreau, 2014). The persistence ratio traces were then averaged across cells.

### Statistics and reproducibility

All experiments were repeated a minimum of three times unless otherwise specified. For comparisons between datasets in Fig. 2E, Fig. 4, Fig. 5, and Fig. 6 the mean of each technical replicate is plotted as a large, colored marker over the single cell distribution. When there were many observations per cell, statistics were done on single cell means. Otherwise, statistics were done on replicate means. Alternatively, in the case of small n, such as Fig. 2G and Fig. 3D, statistics were applied to all measurements. We used a paired t-test when datasets were related (i.e. localization of clathrin and WASP in the same cells (Fig. 1E) or WASP signal at single beads before and after membrane closing (Fig. 3F)) or if there was obvious technical variability (Fig. 4F). We also used paired t-tests to compare migration properties as suggested in (Lord et al., 2020) to reduce influence from day-to-day variability. For measurements that do not exhibit major experimental variation (such as the position of WASP puncta appearance (Fig. 4F)), we used an unpaired t-test. Paired and unpaired two-tailed t-tests are standard for comparing means between parametric datasets.

## Data and code availability

All analysis code and a csv of all data are available at https://github.com/rachel-bot/WASP.

## Online supplemental material

Fig. S1 schematizes the knock-in strategy used and reports the propensity of HL-60 cells to undergo non-homologous end joining and homology directed repair. It also includes sequencing validation of the endogenously tagged WASP and clathrin light chain A cell lines as well as Western blot and sequencing validation of WASP KO cell lines. Fig. S2 shows that endogenous WASP enriches to the interface between HL-60 cells and collagen fibers. Fig. S3 examines the relationship between WASP and FBP17. Fig. S4 demonstrates the resolution increase provided by 3D STED and offers an orthogonal measure of WASP enrichment as a function of bead diameter. Fig. S5 further investigates how polarity informs WASP recruitment. Video 1 shows the dynamics of endogenous WASP in a persistently migrating HL-60 cell. Video 2 shows the spatial separation of endogenous WASP and endogenous clathrin light chain A in an HL-60 cell. Video 3 shows the effect of blocking integrin-based adhesion on WASP puncta dynamics. Video 4 shows the dynamics of WASP recruitment to sites of bead-induced plasma membrane deformation. Video 5 shows plasma membrane and WASP rearrangement as an HL-60 cell migrates over beads. Video 6 shows similar migration between wild type and WASP-null HL-60 cells on flat substrates while Video 7 shows severe defects in WASP-null HL-60 cells plated on nanoridged substrates.

## Abbreviations

Cdc42: cell division control protein 42 homolog
CME: clathrin-mediated endocytosis
DoG: Difference of Gaussians
FBP17: formin binding protein 17
KO: knockout
LCa: light chain A
MSD: mean squared displacement
NPF: nucleation promoting factor
N-WASP: neural WASP
ssDNA: single-stranded DNA
TIRF: total internal reflection fluorescence
WASP: Wiskott-Aldrich syndrome protein
WAVE: WASP family verprolin-homologous protein

## ACKNOWLEDGEMENTS

We thank members of the Weiner lab for conversation and support throughout the project. We acknowledge David Drubin, Sophie Dumont, and Adam Frost for their helpful discussions. We also thank Kirstin Meyer, Suvrajit Saha, Henry de Belly, and Charlotte Nelson for critically reading the manuscript. We thank Pietro de Camilli for providing the anti-FBP17 antibody. This work was supported by NIH F31 HL143882 (RMB), NSF GRFP 1650042 (GRRB), NIH GM118167 (ODW), the NSF Center for Cellular Construction (DBI-1548297), and a Novo Nordisk Foundation grant for the Center for Geometrically Engineered Cellular Systems (NNF17OC0028176).

The authors declare no competing financial interests.

## AUTHOR CONTRIBUTIONS

R. M. Brunetti: conceptualization, data curation, formal analysis, funding acquisition, investigation, methodology, project administration, software, validation, visualization, and writing (original draft and review & editing). G. Kockelkoren: conceptualization, data curation, formal analysis, investigation, methodology, resources, software, visualization, and writing (review & editing). P. Raghavan: investigation, methodology, and resources. G.R.R. Bell: methodology, resources, and writing (review & editing). D. Britain: conceptualization, investigation, and methodology. N. Puri: methodology and resources. S.R. Collins: resources, validation, and writing (review & editing). M. D. Leonetti: conceptualization, methodology, resources, supervision, validation, visualization, and writing (review & editing). D. Stamou: conceptualization, funding acquisition, methodology, resources, supervision, and writing (review & editing). O. D. Weiner: conceptualization, funding acquisition, project administration, supervision, visualization, and writing (original draft and review & editing).

## Supplementary Figures

**Figure S1.**
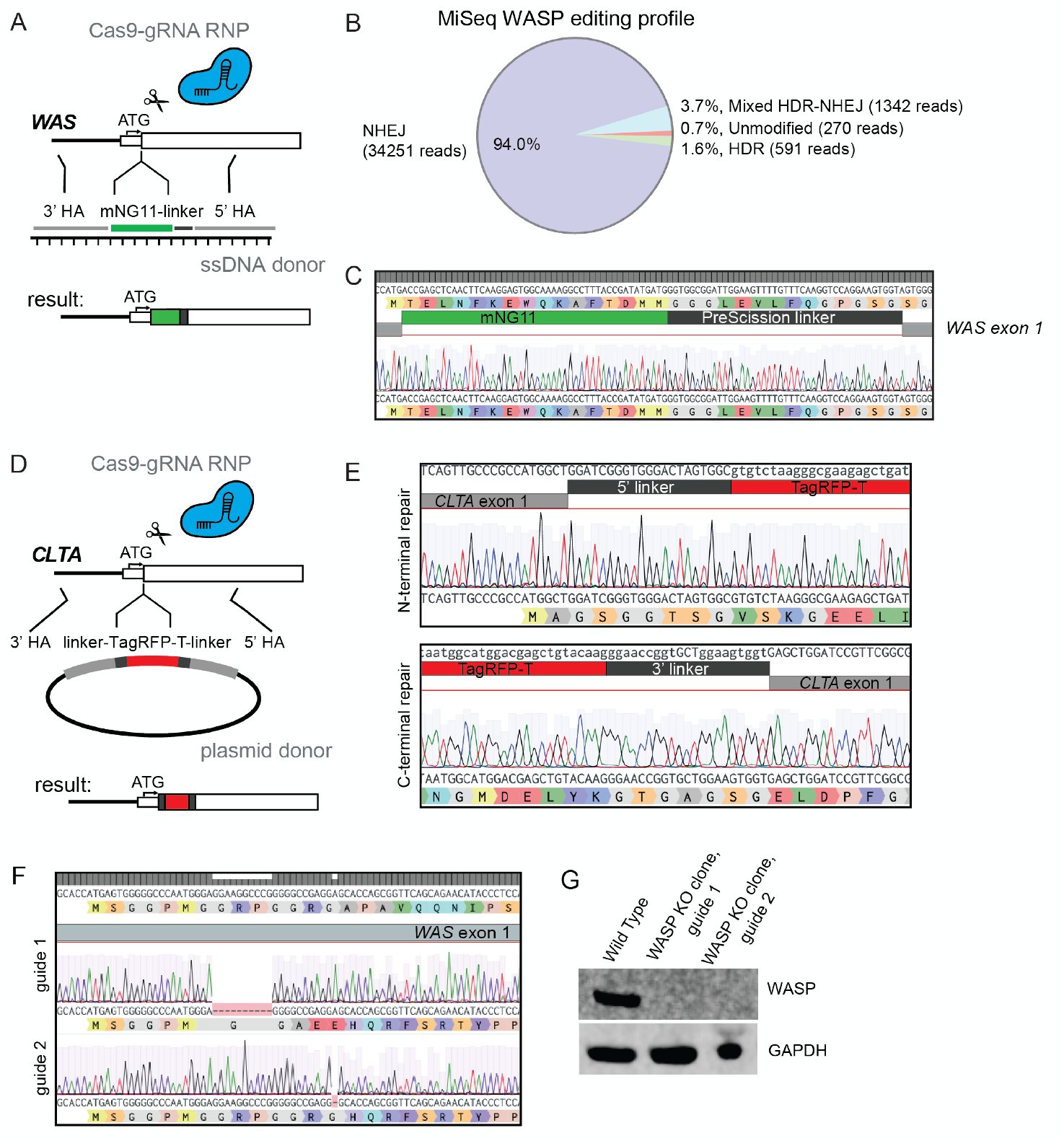
Genetic engineering of HL-60 cells. (A) Strategy for endogenous tagging of WASP with split mNeonGreen2 using a Cas9-sgRNA RNP complex and ssDNA donor. (B) MiSeq results from (A) reveal high cutting efficiency in HL-60s but poor homology directed repair. Analysis of paired end reads was done with CRISPResso2 (Clement et al., 2019). (C) Sequencing of WASP knock-in cells confirms correct insertion of mNeonGreen2_11_ tag. (D) Strategy for endogenous tagging of clathrin light chain A with full length TagRFP-T using a Cas9-sgRNA RNP complex and plasmid donor. (E) Sequencing of an isolated homozygous *CLTA* knock-in clone shows correct repair at both ends of the cut site. (F) Sequence validation of the two clonal WASP KO lines assayed. Both lines have deletions that lead to a frame shift, nonsense, and termination following the end of exon 1. (G) Western blot confirms that WASP is absent in both assayed clonal lines.

**Figure S2.**
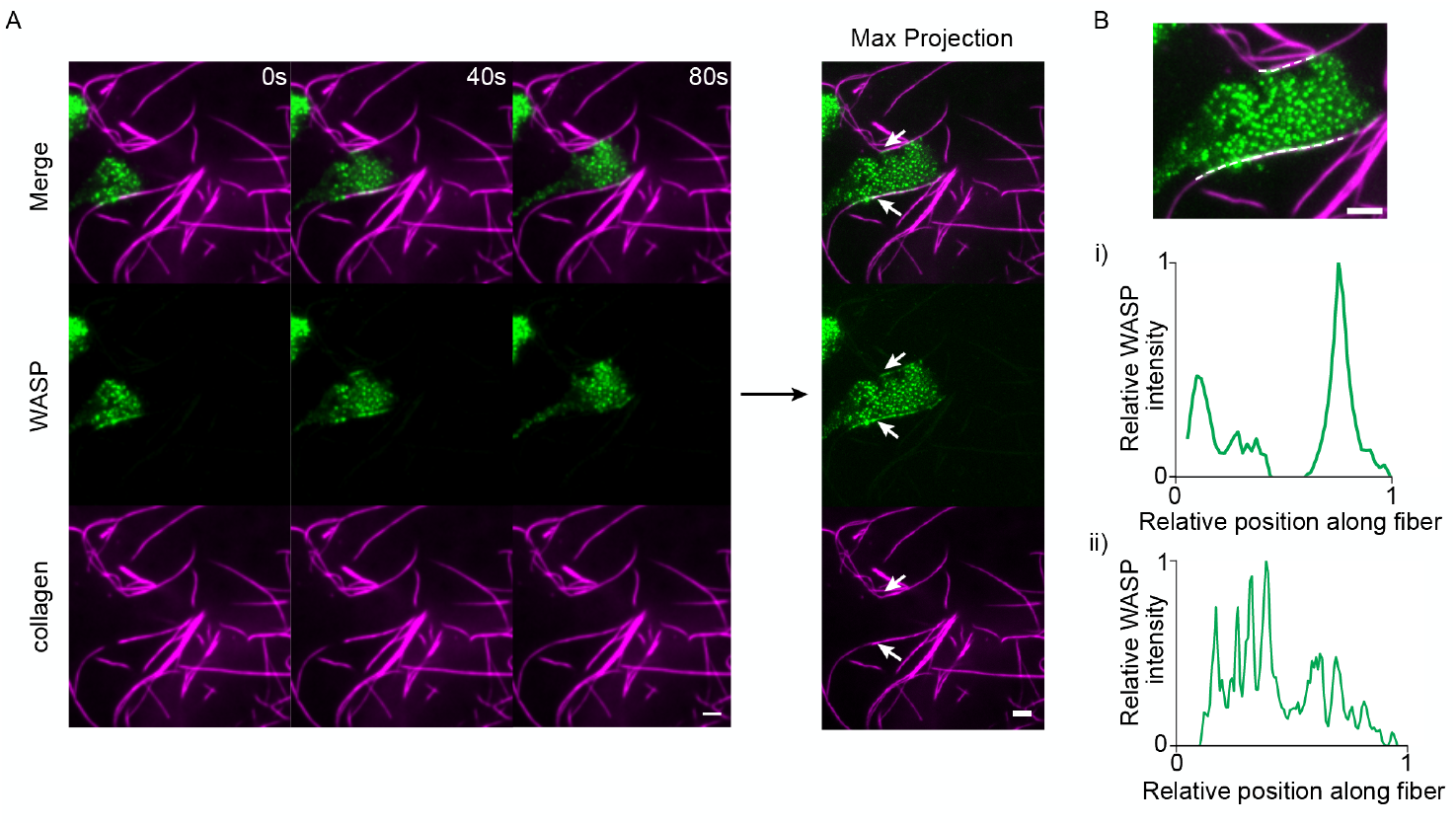
WASP enriches at sites of collagen-induced membrane deformation. (A) Endogenous WASP is recruited to sites of cell (contact with collagen fibers. Right hand side shows the maximum intensity projection over time, which highlights the enrichment of WASP to cell-fiber interfaces. Stretches of contact are marked with arrows. (B) Relative background-subtracted maximum WASP signal along the two fibers marked in (A). (i) is along the upper fiber and (ii) is along the lower fiber, which are marked by dashed lines in the top image. Scale bars are 5 μm. Imaged with TIRF microscopy.

**Figure S3.**
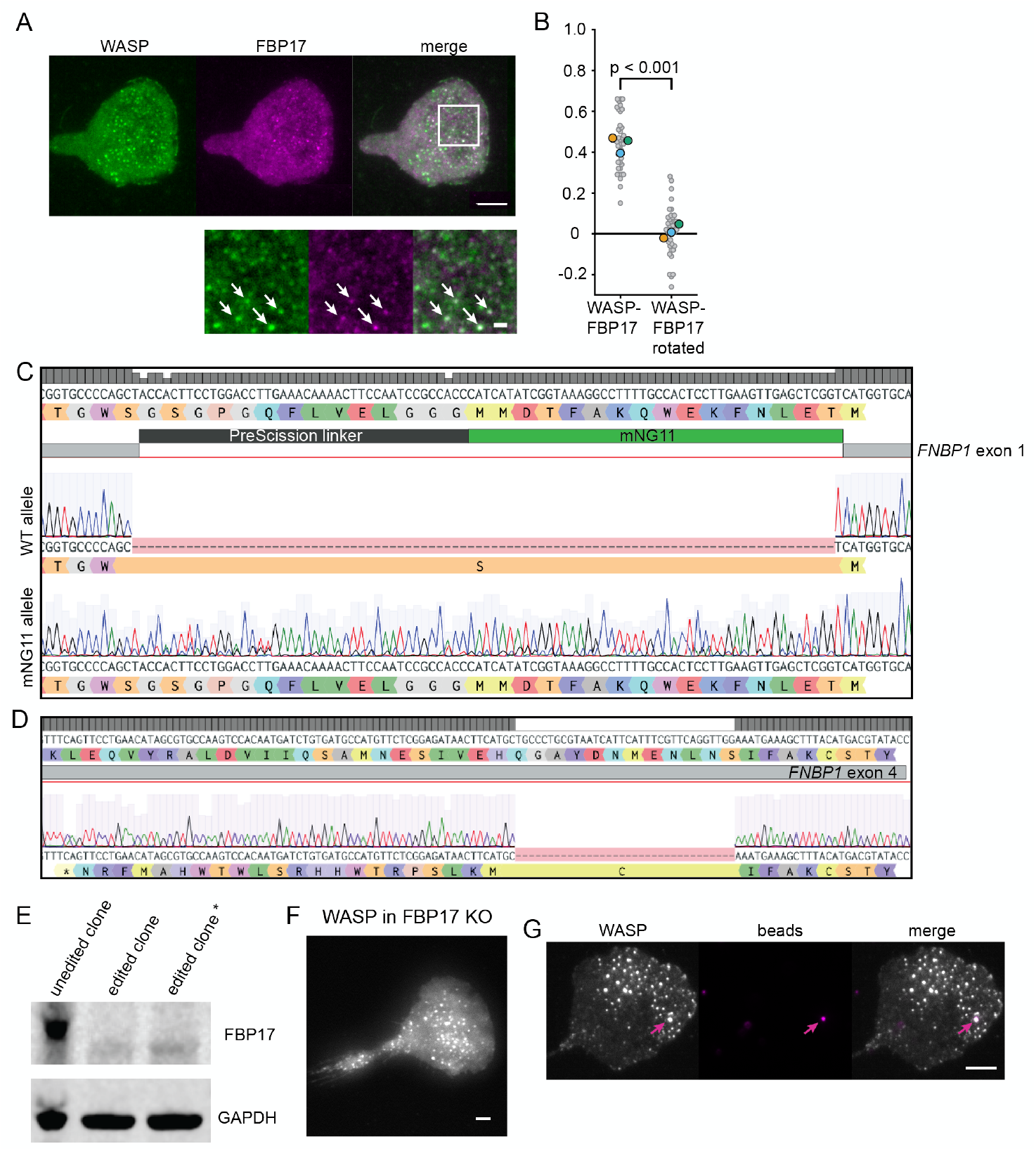
FBP17 co-localizes with WASP but is not required for WASP curvature sensitivity. (A) Overexpressed WASP and endogenous FBP17 co-localize. Below shows a zoomed inset with arrows marking example positions of overlap. Scale bar is 1 μm in the inset. (B) Pearson correlation coefficients between WASP and FBP17 in 38 7.5X7.5 μm ROIs reveal significant co-localization (r = 0.44 ± 0.02) compared to a 90 degree rotated control (0.01 ± 0.02). Data is collected over three experiments. p = 1.51E-17 by a paired two-tailed t-test on the full distribution. (C) Sequencing shows a wild type allele and a correct insertion of mNG2_11_ in the isolated heterozygous FBP17 knock-in clone. (D) Sequencing of an FBP17 KO line shows homozygous deletion of 34 base pairs after amino acid 74, which causes a frame shift and early termination that disrupts the BAR domain (amino acids 1-264). (E) Western blot confirms absence of FBP17 in the assayed FBP17 clonal line (marked with an asterisk) (F) Overexpressed WASP continues to form puncta in an FBP17 KO background. (G) Overexpressed WASP continues to enrich to bead-induced membrane invaginations in the absence of FBP17. Scale bar is 5 μm.

**Figure S4.**
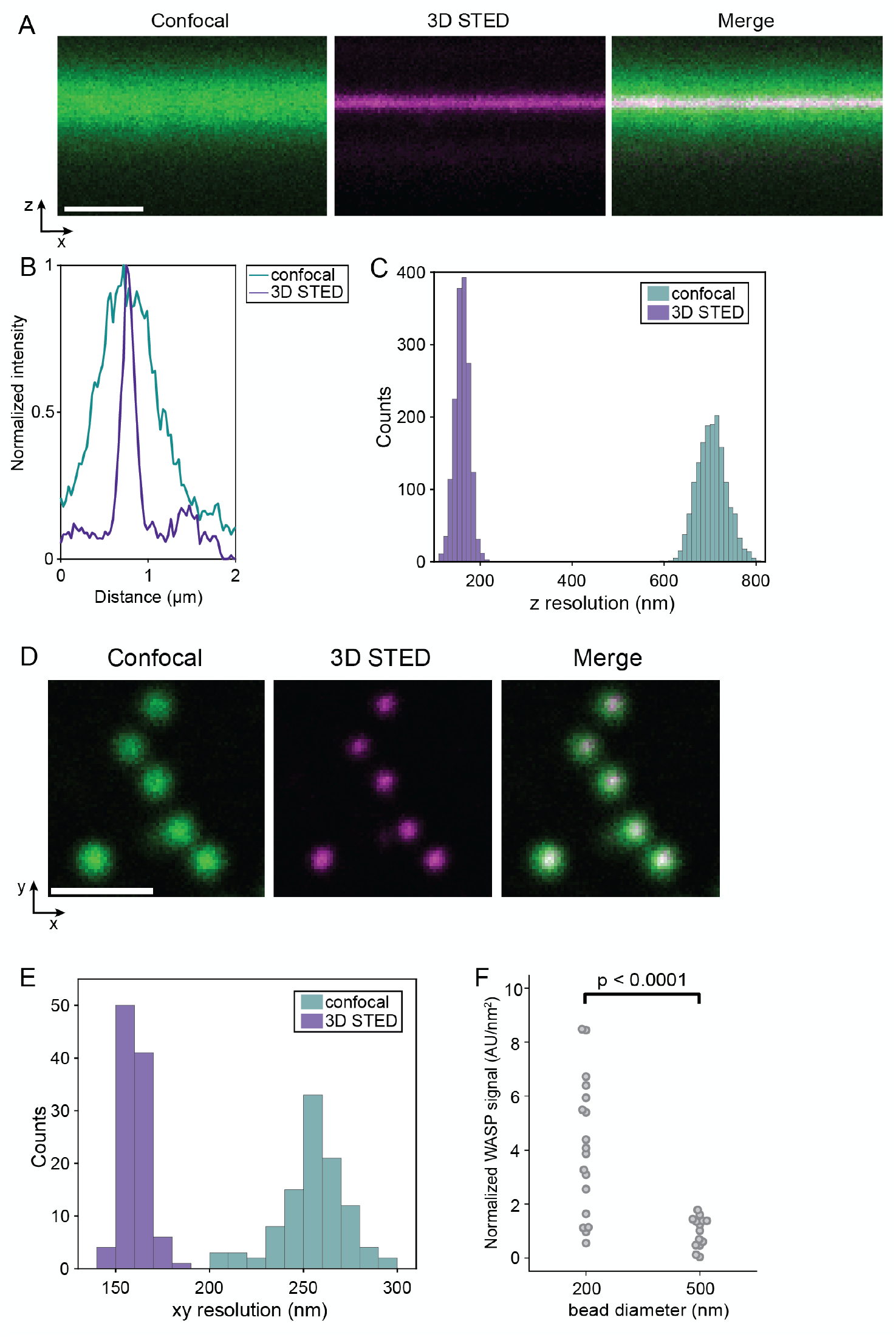
Increased spatial and axial resolution achieved by 3D STED in HL-60 cells. (A) Confocal (left), 3D STED (middle) and merged (right) image of an XZ slice of the ventral membrane of an HL-60 cell labeled with CellMask Deep Red. Scale bar is 1 μm. (B) Representative fluorescence intensity profile along the axial direction in confocal (green) and 3D STED (magenta). (C) Histogram of axial resolution in confocal (green) and 3D STED (purple) imaging mode. 3D STED imaging provides a 4.4x increased axial resolution (160 ± 16 nm for 3D STED versus 704 ± 31 nm for confocal). The axial resolution is determined by the full-width half-maximum of a Gaussian fit to axial intensity profiles of the confocal and 3D STED images (n = 1600 profiles). (D) Estimation of the lateral (XY) spatial resolution offered by 3D STED through imaging fluorescent beads (40 nm) in confocal (left) and 3D STED (right) mode with the same settings as in live-cell experiments. (E) Histogram of spatial resolution in confocal (green) and 3D STED (purple) imaging mode. 3D STED imaging provides a 1.6x increased spatial resolution (160 ± 6 nm for 3D STED versus 255 ± 17 nm for confocal). Spatial resolution is calculated by the full-width half-maximum of 2D Gaussian fits to the beads (n = 102 beads). Scale bar is 1 μm. (F) Integrated intensity of thresholded WASP signal across XYZ stacks reveals significant enrichment of WASP to 200 nm beads compared to 500 nm beads (non-neck invaginations) when normalized to bead surface area. Mean normalized WASP intensity per unit area is 4.08 ± 0.59 AU/nm^2^ for 200 nm beads and 0.97 ± 0.15 AU/nm^2^ for 500 nm beads. p = 8.81E-05 by an unpaired two-tailed t-test on the normalized WASP intensities at all beads. n_200nm_ = 18 beads collected from two experiments and n_500nm_ = 14 beads collected from one experiment.

**Figure S5.**
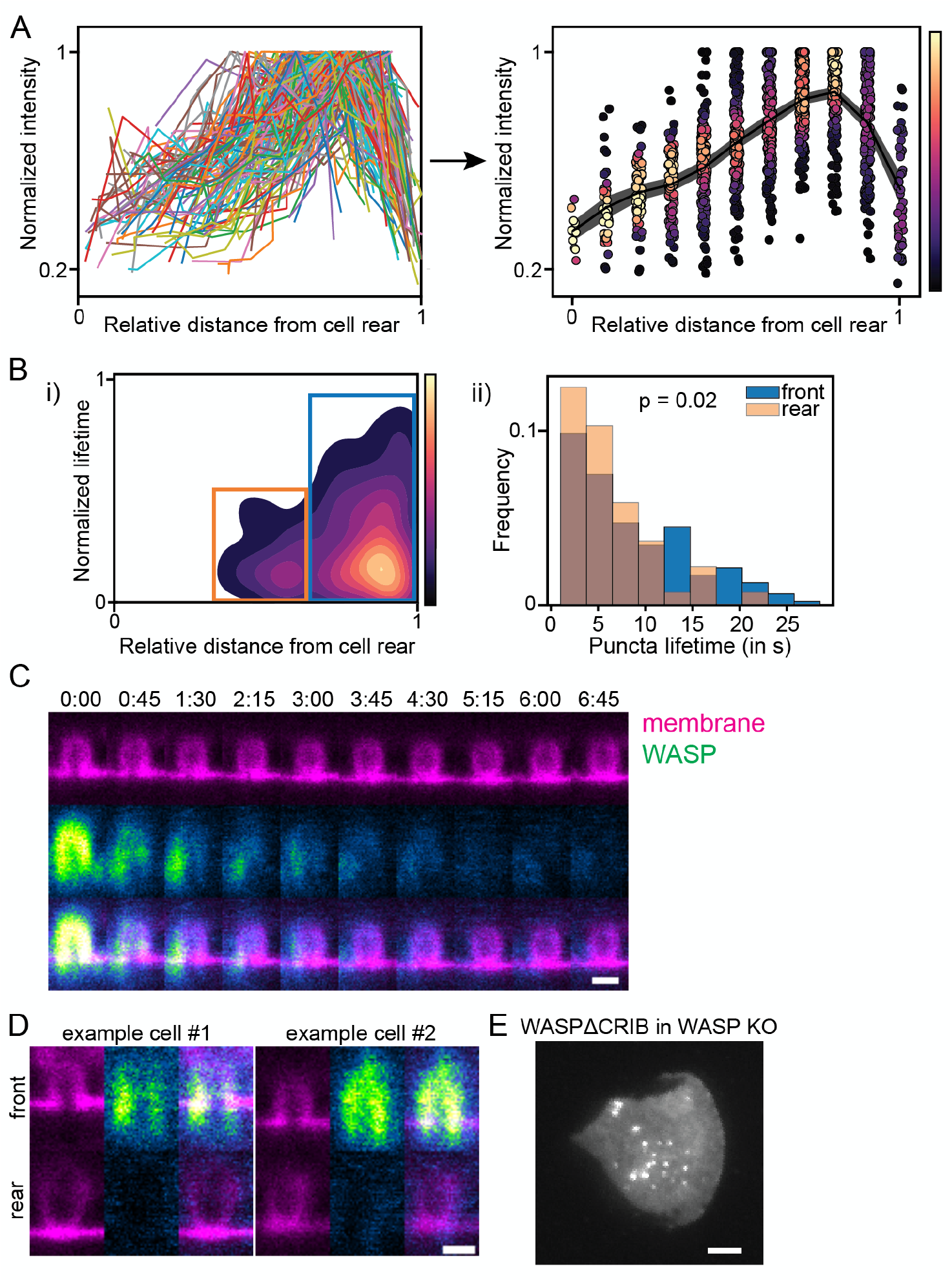
Further investigation of the effect of polarity on WASP recruitment and dynamics. (A) Normalized traces of integrated WASP signal at 135 beads that travel the cell length. Data was collected across three experiments. Traces exhibit consistent behavior, peaking in the front half of the cell and falling off as the bead approaches the cell rear. Right side of the panel shows the averaged data with individual points overlaid and colored by data density. (B) (i) Kernel density estimation comparing the position of puncta appearance with lifetime reveals two populations (boxed). (ii) Histograms of the boxed populations show puncta that appear closer to the cell rear extinguish faster, suggesting that position within the cell influences disappearance. More than 200 puncta were collected across 7 cells from two experiments. Mean lifetime is 8.50 ± 0.49 s for puncta that nucleate in the front 35% of the cell (blue box) and 6.24 ± 0.67 s for puncta that nucleate further back (orange box). p = 0.02 by an unpaired two-tailed t-test on all puncta lifetimes. (C) Time lapse STED imaging shows membrane invaginations maintain geometry but lose WASP signal as the cell continues moving. Scale bar is 500 nm. (D) Single XZ STED slices of beads at the cell front and cell rear of the same cell show that despite similar geometry WASP is only recruited to invaginations at the front. Images are scaled to the same intensity. Scale bar is 500 nm. (E) WASP KO cells rescued with endogenous levels of fluorescent WASPΔCRIB form puncta. Scale bar is 5 μm.

## Video Legends

**Video 1. Distribution of endogenous WASP-mNeonGreen2 in a persistently migrating HL-60 cell (as in Fig. 1A).** WASP enriches to foci on the ventral surface of the cell that are concentrated toward the cell front. Imaged with TIRF microscopy and displayed with an inverted color map. Scale bar is 5 μm. Time, mm:ss.

**Video 2. Endogenous WASP-mNeonGreen2 and endogenous clathrin light chain A-TagRFP-T remain spatially separate in a migrating HL-60 cell (as in Fig. 1D).** Scale bar is 5 μm. Imaged with TIRF microscopy. Time, mm:ss.

**Video 3. WASP forms puncta that are stationary relative to the substrate but undergo retrograde flux in the absence of integrin engagement (+EDTA) (as in Fig. 1F).** Scale bar is 5 μm. Imaged with TIRF microscopy.

**Video 4. Endogenous WASP (gray) is recruited to all 200 nm beads (magenta) as the cell encounters them (as in Fig. 2C).** WASP signal is sustained over the cell front but begins to diminish as beads reach the cell rear. Scale bar is 5 μm. Imaged with TIRF microscopy. Time, mm:ss.

**Video 5. Evolution of the membrane and WASP as confined cells run over beads (2D counterpart to Fig. 3C).** With time, the membrane closes around the bottom of the bead, transitioning from an open hole to a sub-resolution closed neck. At the same time, WASP redistributes from coating the open membrane hole to focal enrichment at the closed neck. Arrows mark examples where this occurs. WASP is imaged with confocal microscopy and the membrane is imaged with 2D STED microscopy. Scale bar is 1 μm. Time, mm:ss.

**Video 6. WASP KO HL-60 cells exhibit relatively minor migration defects on flat substrates (as in Fig. 6C and G-I, top).** Cells are labeled with NucBlue for tracking and imaged for one hour at 37°/5% CO2. Overlaid are cells trajectories that last at least half the movie. Scale bar is 10 μm. Time, mm:ss.

**Video 7. WASP KO HL-60 cells exhibit significant migration defects on textured (800 nm nanoridged) substrates (as in Fig. 6D-F and G-I, bottom).** Cells are labeled with NucBlue for tracking and imaged for one hour at 37°/5% CO2. Overlaid are cell trajectories that last at least half the movie. Scale bar is 10 μm. Time, mm:ss.

